# An exploratory data analysis from ovine and bovine RNA-seq identifies pathways and key genes related to cervical dilatation

**DOI:** 10.1101/2023.02.07.526593

**Authors:** Joedson Dantas Gonçalves, José Bento Sterman Ferraz, Flávio Vieira Meirelles, Ricardo Perecin Nociti, Maria Emilia Franco Oliveira

## Abstract

The present study developed a review and exploration of data in public and already validated repositories. The main objective is to identify the pathways involved in ruminant’s cervical dilatation, which are conserved between cattle and sheep in the follicular and luteal phases of the reproductive cycle. In cattle, 1961 genes were found to be more expressed in the follicular phase and 1560 in the luteal phase. 24 genes were considered exclusively expressed from these 18 genes were in the follicular phase and 6 genes were in the luteal phase. In sheep, 2126 genes are more expressed in the follicular phase and 2469 genes are more expressed in the luteal phase. Hoxb genes were identified in both species and are correlated with the PI3K/Akt pathway. PI3K/Akt was also found in both cattle and sheep, appearing prominently in the follicular and luteal phases of both species. Our analyzes have pointed out that the PI3K/Akt pathway and the Hoxb genes appear in prominence, in modulating mechanisms that involve estrus alterations in the cervix. PI3K/Akt appears to be an important pathway in the cervical relaxation process.

## 1. Introduction

The anatomy of the ovine cervix is one of the main limiting factors for cervical transposition in sheep, due to the number, internal diameter, and distribution of cervical rings [1]. Even with cervical remodeling during the estrus phase, cervical penetration for artificial insemination procedures in ewes remains problematic [2,3]. The challenge is even greater during the luteal phase when embryo collection is performed [4]. A series of studies attempted to develop protocols with satisfactory cervical relaxation responses, for better application of reproductive biotechnologies [3]. The mechanism of cervical dilatation in sheep is complex, involving several substances, proteins, enzymes, and hormones [4].

In ruminants, both at parturition and estrus, increased estrogen concentration appears to initiate a cascade of events that culminate in cervical relaxation [5]. There is a major obstacle to unraveling the mechanisms of cervical ripening, as the mechanisms in mammals are highly variable at parturition, which is the time of greatest dilatation. A practical example would be that in humans and primates, the placenta produces large amounts of progesterone during pregnancy and childbirth. Most other placental mammals, such as sheep, mice, rabbits, and rats, have systemic progesterone withdrawal, in which the serum concentration of progesterone decreases before labor begins [6].

The enzyme 5-α-steroid reductase (SRD5A1) is essential in cervical regulation and remodeling, and it has been observed that SRD5A1 mRNA expression is found only in mice, suggesting that this well-documented mechanism of cervical ripening is specific to mice. Although, was found in guinea pigs, which belong to a rodent lineage basal branching [7]. It has also been reported that the enzyme HSD17B1, known mainly for its role in the synthesis of estradiol, is not expressed in the cervix of the opossum and rat, little expressed in the cervix of the armadillo, highly expressed in guinea pigs at all stages of the reproductive cycle, and in rabbits at the end of pregnancy [6]. Moreover, these are suggestive that in species with common ancestors, there are conserved cervical dilatation mechanisms, as well as variations of these mechanisms among species.

Furthermore, addressing those topics a review and data mining were carried out in public and already validated repositories. We hypothesized that differentially expressed genes and divergent biological processes would indicate differences and similarities related to cervical relaxation in cows and sheep. The main objective was to identify the pathways involved in cervical dilatation, which are conserved between cattle and sheep in the follicular and estrous phases of the reproductive cycle. Such results are important to better understand the mechanism of cervical dilatation in sheep, and indicating signaling pathways, which may help to understand and improve the efficiency of cervical relaxation protocols. Thus, consequently, improve the application of reproductive biotechniques in ruminant species.

## 2. Results

In cattle, 1961 genes were found to be more expressed (padj<0.1 and |log2foldchange|>0.6) in the follicular phase and 1560 in the luteal phase. A total of 24 genes were considered exclusive of these 18 genes in the follicular phase and 6 genes in the luteal phase. In sheep, 4595 genes were differently expressed (padj<0.1 and |log2foldchange|>0.6) out of a total of 19581 expressed genes. Being 2126 more expressed in the follicular phase and 2469 genes more expressed in the luteal phase. 4 unique genes were found in the follicular phase.

The signaling pathways found have different functions and expression intensities according to the phase of the estrous cycle. Signaling pathways are linked to genes of different categories: adrenergic, dopaminergic, purinergic receptors, growth factors, tumorigenesis, hematopoiesis, chromatin regulation and condensation, transcriptional regulation, mucins, hormone ligands, cartilage and genes involved in vasoconstriction, vasodilation, and muscle contraction.

Figure 1 presents a summary of the analysis of differences in gene expression in cattle in the follicular and luteal phases. The differently expressed genes can be visualized in a volcano plot in Figure 1A, with the genes in red in the follicular phase and the genes in blue in the luteal phase. In the follicular phase, ENSBTAG00000048276 (trefoil factor 1), TMPRSS11B N-terminal-like (TMPRSS11BNL), LOC112441508 and BPI fold containing family A, member 2B (BPIFA2B) were more expressed. In the luteal phase, the most differently expressed genes were ENSBTAG00000011470, transmembrane inner ear (TMIE), teratocarcinoma-derived growth factor 1 (TDGF1), LDL receptor related protein 2 (LRP2), solute carrier family 30 member 8 (SLC30A8). In Figure 1B we have the results of the PCA analysis, demonstrating the clustering of data. Figure 1C shows 30 different pathways that are modulated by these genes, with the PI3K/Akt pathway having the highest number of genes expressed. In Figure 1D we have the network of genes with different expressions in which the interpellation of genes with the other pathways can be observed. We can observe the PI3K/Akt pathway in the center of the network.

**Figure 1.**
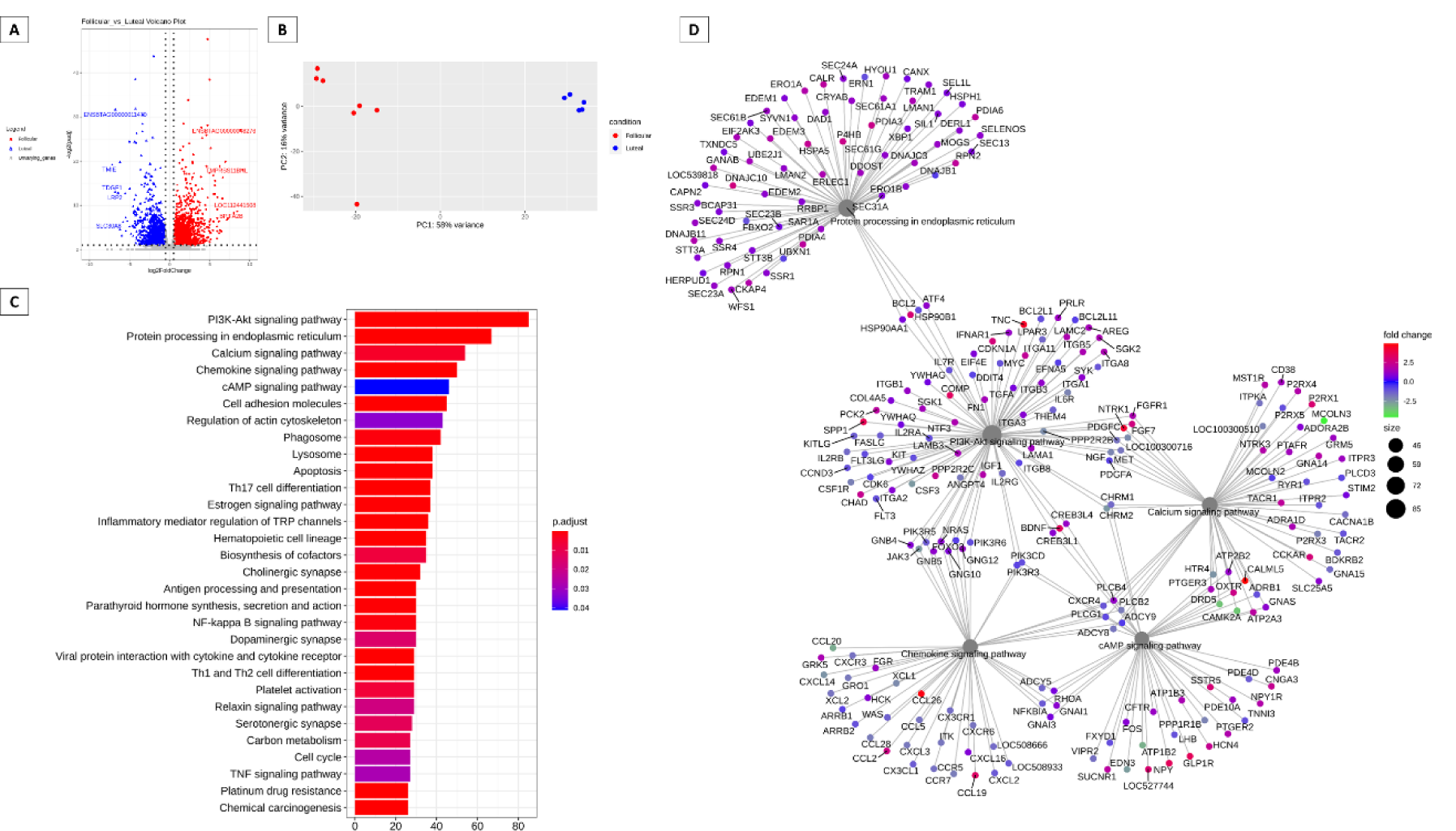
Summary of the different gene expression analysis in bovine cervix during the follicular and estrous phases. (A) Volcano-plot comparing follicular and estrous phases. (B) PCA analysis. (C) Barplot KEGG enrichment pathways analysis of differentially expressed genes (D) Network analysis based on the of pathways and differentially expressed genes during the follicular and luteal phase of bovine cervix.

Figure 2 presents the analysis of differences in gene expression in sheep in the follicular and luteal phases. The differently expressed genes can be visualized in the volcano plot in Figure 2A, with the genes in red in the follicular phase and the genes in blue in the luteal phase. In the follicular phase, Dopamine Receptor D2 (DRD2), ENSOARG00020011601, Secreted LY6/PLAUR domain containing 1 (SLURP1), ENSOARG00020000537, Keratinocyte differentiation-associated protein (KRTDAP) were differently expressed. In the luteal phase, the most expressed genes were ENSOARG00020024493, ADAM metallopeptidase domain 7 (ADAM7), keratin, type II microfibrillar, component 5-like (KRT85), ENSOARG00020010220, ENSOARG00020015599. In Figure 2B we have the results of the PCA analysis, demonstrating the clustering of data. Figure 2C shows 30 different pathways that are modulated by these genes, with the Neuroactive ligand-receptor interaction pathway having the highest number of genes expressed, followed by the PI3K/Akt pathway. In Figure 1D we have the network of genes with different expressions in which the interrelationship of genes with the other pathways can be observed. We can also observe the PI3K/Akt pathway near the center of the network and peripherally the signaling pathways of Ras and RAP1.

**Figure 2.**
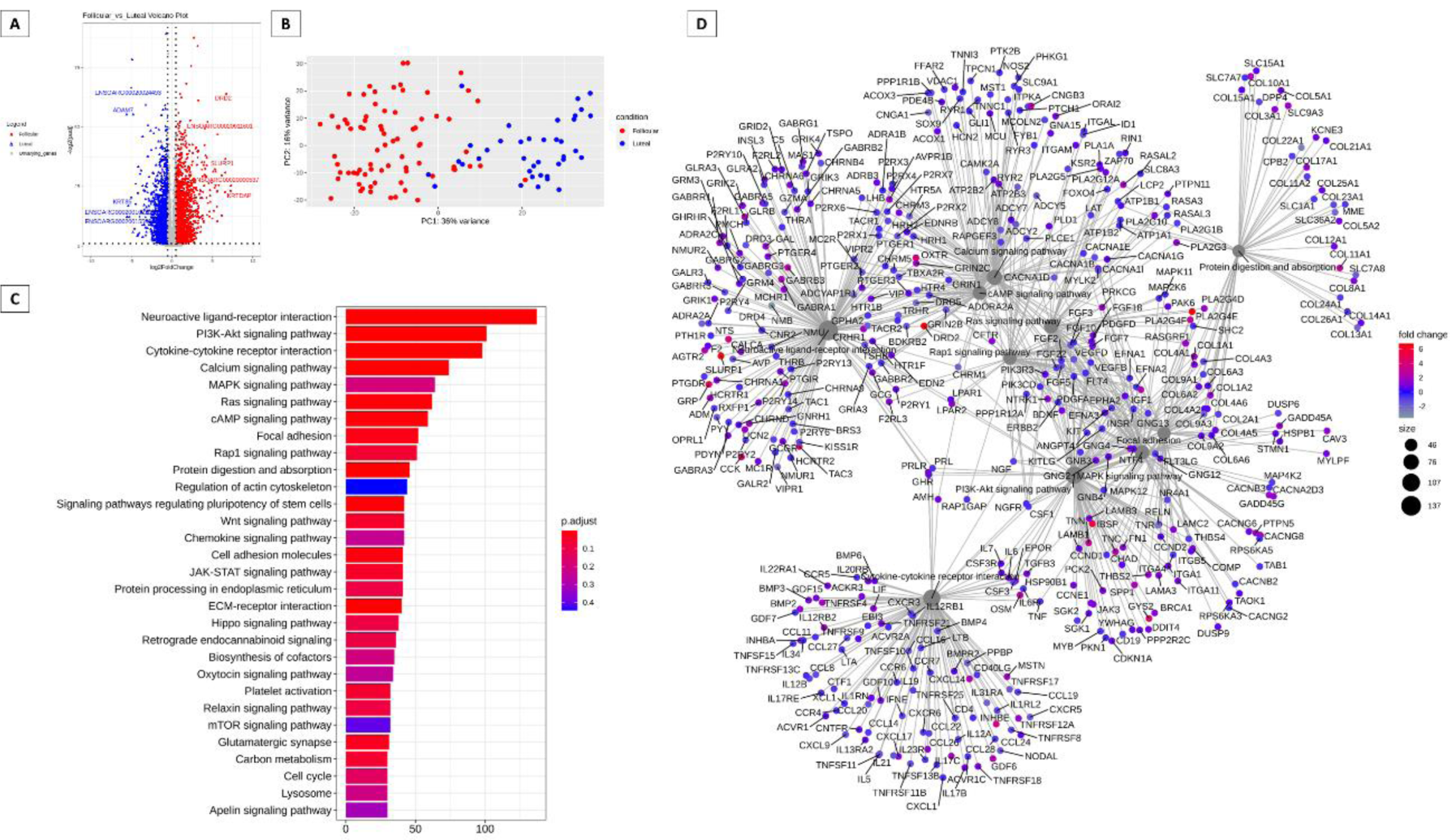
Summary of the analysis of the difference in gene expression of the ovine cervix in the follicular versus luteal phase. (A) Volcano plot of RNA sequencing (RNAseq) comparing follicular and estrous phases. (B) PCA analysis. (C) Network analysis based on the Kyoto Encyclopedia of Genes and Genomes [KEGG] enrichment of differentially expressed genes (D) Enriched network of pathways and genes expressed in the follicular and luteal phase of the ovine cervix.

Figure 3 summarizes the modulation of key genes and their possible targets in the bovine cervix (Figure 3A). These pathways may be modulated by key genes. The key gene is considered the highest degree of connectivity to other genes to promote a molecular event [8]. In the interaction network Figure 3B, again the PI3K/Akt pathway appears to be modulated. In Figure 4 we have the pathways related to the key genes in sheep. In Figure 4 A we have the main pathways related to key genes and the network of pathways with PI3K/Akt in the center, also modulated by these key genes (Figure 4B).

**Figure 3.**
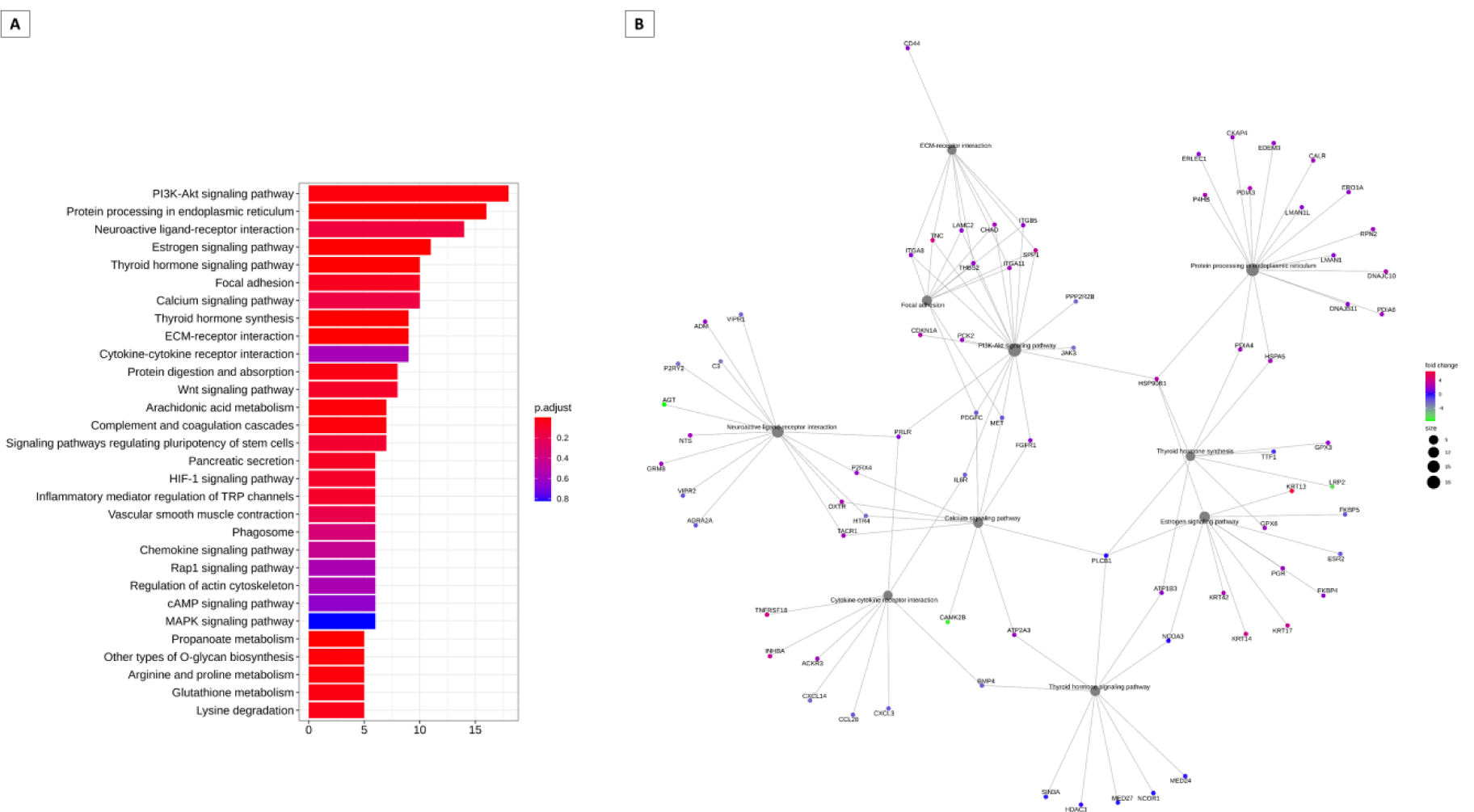
Analysis of enrichment of KEGG pathways that can be regulated by key transcription factors in cattle. (A) Main gene pathways with interaction with key genes. (B) Analysis of gene networks with interaction with key genes.

**Figure 4.**
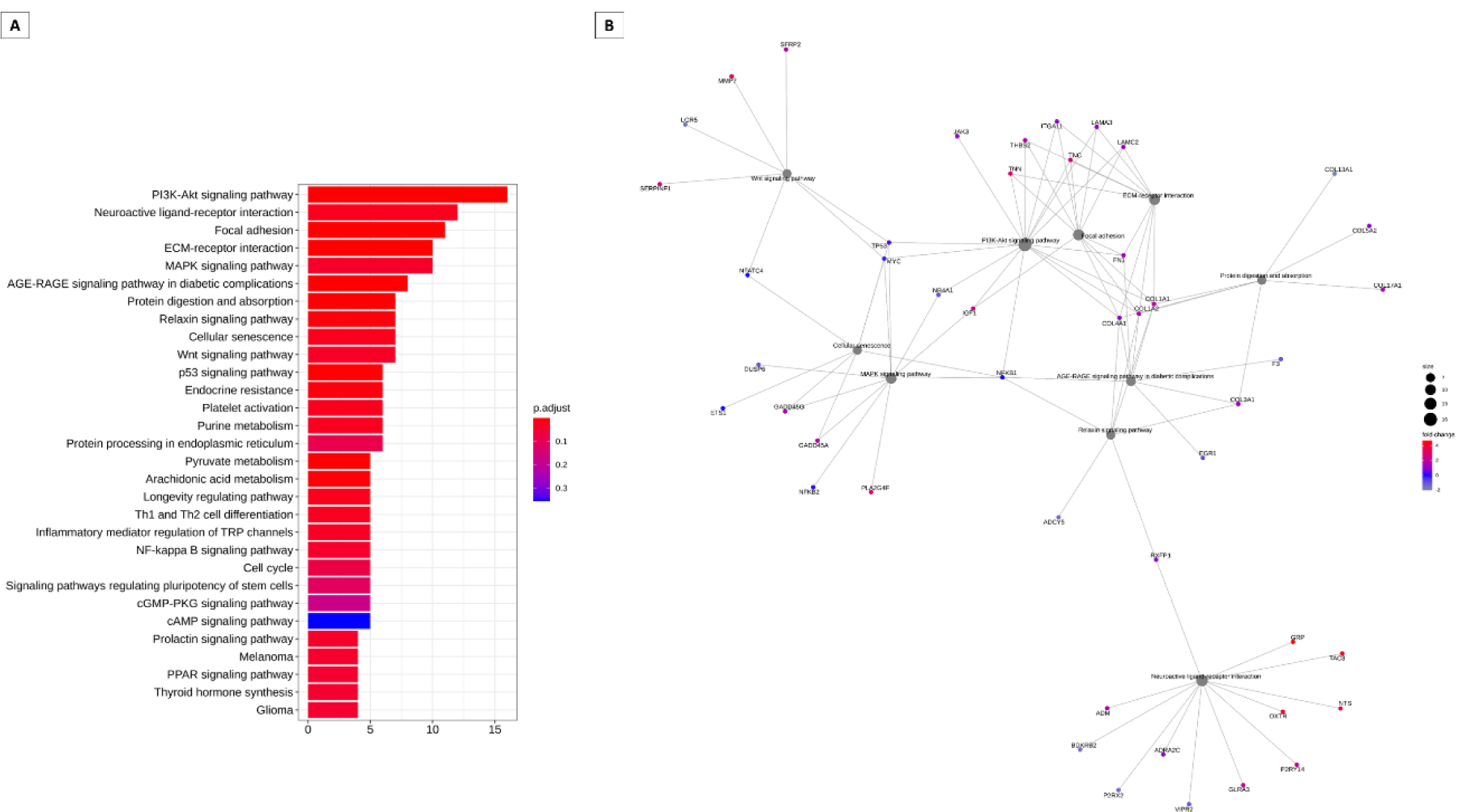
Analysis of enrichment of KEGG pathways that can be regulated by key transcription factors in sheep. (A) Main gene pathways with interaction with key genes. (B) Analysis of gene networks with interaction with key genes.

In Figure 5, pathways and genes can be modulated by Hoxb genes in cattle (Figure 5) and sheep (Figure 6). Hox genes are essential in directing the further development of tissues in the embryonic stage [9]. It can be observed that the PI3K/Akt pathway appears to be modulated by these Hox genes, both in the follicular and luteal phases, appearing prominently in the interaction networks. It is also observed in the interaction networks, that several other genes may also be modulated by the Hoxb genes. These results were obtained using Spearman’s correlation with a P value < 0.05 and an absolute R-value greater than 0.5.

**Figure 5.**
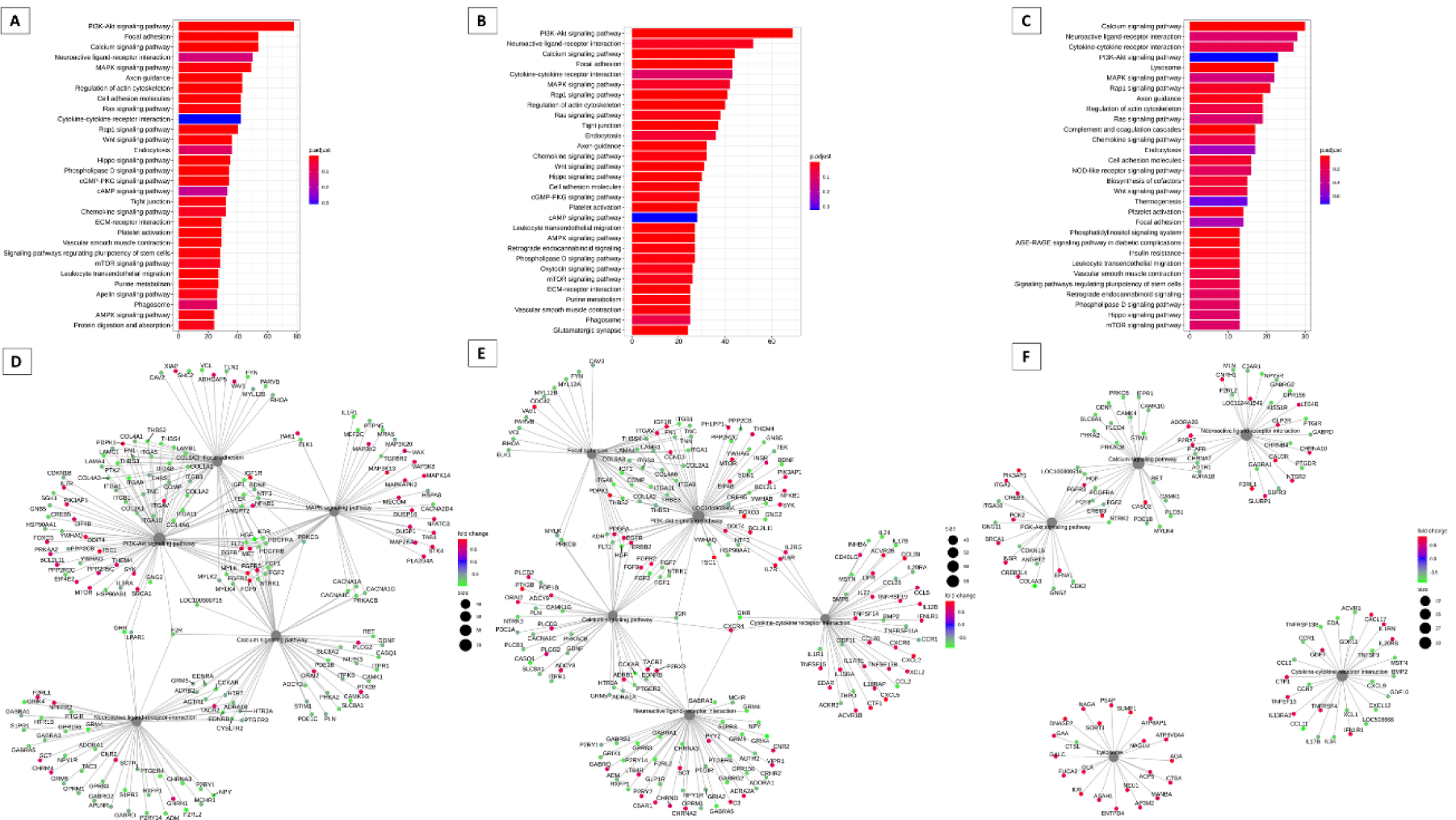
Analysis of enrichment of KEGG pathways that can be regulated by Hox genes in cattle. (A)-Hoxb3, (B)-Hoxb8, (C)-Hoxb9, (D)-Hoxb3, (E)-Hoxb8, (F)-Hoxb9.

**Figure 6.**
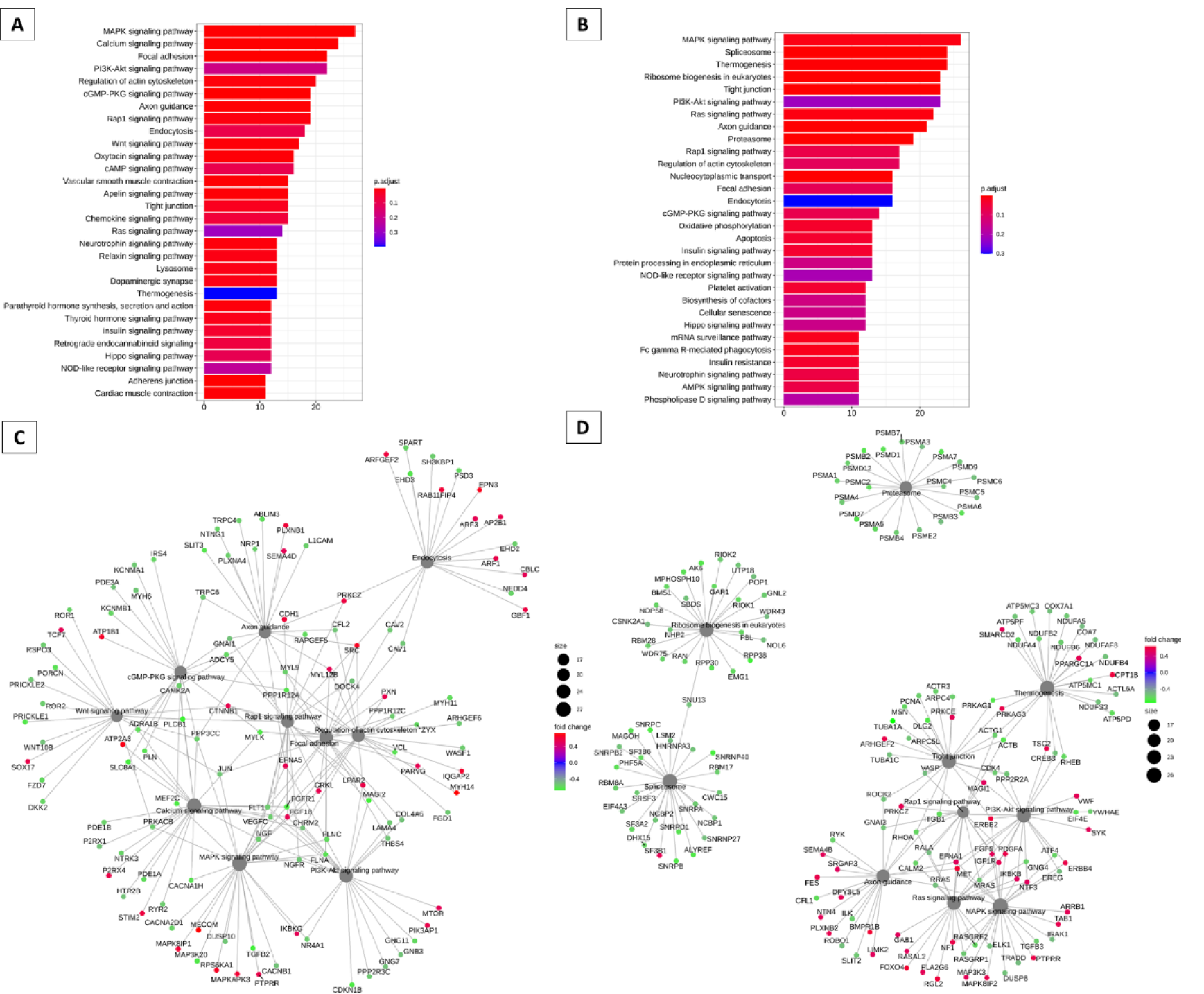
Analysis of enrichment of KEGG pathways that can be regulated by Hox genes in sheep. (A)-Hoxb2, (B)-Hoxb3, (C)-Hoxb2, (D)-Hoxb3.

In the supplementary material, the hub genes most expressed in cattle (Table S1) and sheep (Table S2) are demonstrated. In the follicular (Positive Log Fold Change) and luteal (Negative Log Fold Change) phases. As a hub gene most expressed in the follicular phase in cattle, mucin 1 (MUC1) (Log Fold Change 1.5759491672159) has a fundamental role in the cervical mechanism. For sheep, the hub genes most expressed in the cervix during the follicular and luteal phases are shown in Table S2. The hub gene most expressed in the follicular phase of sheep was Estrogen receptor 1 (ESR1) (Log Fold Change of 1.01079315236001) and Estrogen receptor 1 (ESR2) was highly expressed in the luteal phase (Log Fold Change of −1.10402057614656).

Table S3 shows the 100 genes most expressed in the follicular and luteal phase of cattle found in the evaluation. The BPI fold containing family A, member 2ª (BPIFA2A) was highly expressed (Log Fold Change of 10.4876703575293) in the follicular phase. Well expressed in the luteal phase in cattle Table S3, angiotensinogen (AGT) (Log Fold Change of −7.72323486386274), a participant in the renin-angiotensin system (RAS). The 100 most expressed genes in the follicular and luteal phase of the sheep cervix are shown in Table S4. Found in the follicular phase, Trefoil factor 1 (TFF1) (Log Fold Change 5.31535865509055), while in the luteal phase, ADAM metallopeptity domain 7 (ADAM7) was the most expressed gene (Log Fold Change 5.58250234721926).

The genes considered exclusive in the bovine cervix in the follicular and luteal phases are presented in Table S5. 23 genes were found, of these 17 genes were in the follicular phase (Positive Log Fold Change) and 6 genes were in the luteal phase (Negative Log Fold Change). BPI (BPIFA2B) was the most expressed in the follicular phase of the bovine cervix (Log Fold Change of 8.01169488659356) and Retinol dehydrogenase 16 (RDH16) (Log Fold Change of −5.30278893747946) most expressed in the luteal phase.

## 3. Discussion

The cervix is a complex fibrous structure that undergoes structural changes according to the estrous cycle phase. In the follicular phase, the cervix is more open for the reception and transport of sperm [1]. In the luteal phase, the cervical canal is completely closed to protect against pathogenic microorganisms [10]. There is a biotechnical interest in cervical dilatation during the luteal phase in sheep due to the difficulty of transposition of the cervix. There is, however, more information related to cattle in the literature. Thus, a comparison of the mechanisms between sheep and cattle may be an alternative for a better understanding of the cervical relaxation event. Our findings point to molecular differences in the cervical physiology of cattle and sheep.

In the bovine follicular phase, 4 genes were found differently expressed: TFF1, TMPRSS11BNL, LOC112441508, and BPIFA2B. TFF1 is encoded by an estrogen-responsive gene [11,12]. There are reports that the synthesis of secretory mucins is typically accompanied by the co-secretion of TTF peptides [13]. TFF peptides help maintain the surface integrity of mucosal epithelia [14]. TFF1 expression has been reported in small amounts in the human endocervix [15]. These triphilic peptides are involved in the protection and restoration of epithelia [15] in addition to binding to mucins [16,17] and mucin-associated proteins [18] In the follicular phase, there is a large amount of cervical mucus production. This cervical mucus helps in cervical dilation, due to the release of substances that bind to collagen and cause distension of the fibers [19]. TMPRSS11B, also found in the follicular phase, is a member of the type II trypsin-like serine protease [20], expressed on the cell surface [21]. There are reports of data from the immunohistochemical analysis that show that TMPRSS11B is expressed in tissues of some types of cancer [21]. Serine proteases are related to the expression of pro-inflammatory cytokines [22]. The inflammatory process is important in the dilation mechanism, as it recruits defense cells. The neutrophil, for example, seems to be an important component for cervical softening, as the collagenase released by neutrophils is very important for the disruption of collagen fibers, which are the main structural elements of the cervix. This disruption causes distention of the organ [23].

BPIFA2B, highly expressed in the follicular phase of the bovine cervix, is found in the secretory granules of neutrophils and is secreted in response to activation of Toll-like receptor (TLR)-mediated signaling, where it acts as an innate immune effector protein permeabilizing the plasma membrane of Gram negative bacteria [24–26]. In estrus, the cervix is open under the effect of estrogen, so the open channel allows the entry of bacteria from the normal flora of the vagina into the uterine lumen [1]. Thus, BPIFA2B may have its role related to antimicrobial activity. This gene is also considered exclusive in the bovine cervix in the estrous phase (Table S5).

In the luteal phase in cattle, the 5 differently expressed genes were ENSBTAG00000011470, TMIE, TDGF1, LRP2, SLC30A8. The TMIE already reported in adult mice and rats is expressed in various tissues [27]. However, the role of TMIE is still uncertain, there are only conclusions of alterations in mice with TMIE deficit in the inner ear, as this gene is involved in sensory mechanotransduction in cochlear hair cells [28]. In other tissues, whether in humans, rats, or other mammals, there are no reports of the functions of TMIE. STAT3 protein inhibitor has already been identified as a potential binding partner [29]. STAT3 is already reported in the remodeling of stroma and uterine epithelium in the luteal phase, mainly during embryonic implantation [30]. Thus, the TMIE may be involved in the Mechanotransduction of signals involving structural changes in the cervix.

TDGF1 is a member of the epidermal growth factor (EGF) - Crypto-1/fibroblast growth factor (FRL1) related ligand and acts as a ligand for activation of the src-Akt pathway [31] and is involved in embryogenesis and tumorigenesis [32–34]. Its participation is essential for early embryonic development and maintains the pluripotency of embryonic stem cells in mice and cattle [35,36]. Secreted soluble forms of TDGF1 may also activate the PI3K/Akt pathway [37]. Similar to other members of the EGF family, it can stimulate cell proliferation, migration, angiogenesis, invasion, cell survival, and epithelial-mesenchymal transition [38,39]. Furthermore, this gene is totally related to cell stimulation, growth, and differentiation, its action may be linked to cervix physical changes during the estrous cycle. Another gene highly expressed in the luteal phase in cattle is LEP2, an endocytic receptor strongly expressed in steroid-responsive tissue and epithelial cell types, particularly in male and female reproductive organs such as the epididymis, prostate, ovaries, and uterus [40]. It may also have effects on cervical mechanical characteristics because its action on female reproductive systems is related to the alteration of uterine architecture, being positively regulated by P4.

In the ovine follicular phase, the 5 genes were differently expressed, DRD2, ENSOARG00020011601, SLURP1, ENSOARG00020000537, KRTDAP. DRD2 is considered a dopamine receptor with a role in motor control and neuroendocrine activities [41]. In male reproductive organs, DRD2 has been linked to the stimulation of penile erection, mediated by the oxytocin pathway [42,43]. In females, oxytocin is responsible for the excitability and contractility of the uterus [44], in this way, DRD2 as a dopaminergic receptor can act directly or indirectly for these stimuli to occur. SLURP1 facilitates the functional development of T cells and suppresses the production of TNF-a (Tumor Necrosis Factor-alpha) by T cells, secretion of IL-1 b and IL-6 by macrophages, and in humans the secretion of IL-8 by the intestine [45]. This affinity for interleukins and tumor necrosis factor may be linked to the mechanism of cervical dilatation, as these interleukins actively act on the cellular remodeling of the cervix and, consequently, on cervical relaxation [46,47]. These two genes are also highlighted among the 100 genes most expressed in the follicular phase of sheep (Table S4).

In the luteal phase, the most differently expressed genes were ENSOARG00020024493, ADAM7, KRT85, ENSOARG00020010220, ENSOARG00020015599. More recently, it was reported that ADAM7 overexpression strongly promoted cell proliferation, migration, and invasion and inhibited cell apoptosis in trophoblastic cells [48]. In addition, there are reports that matrix metalloproteinases, such as ADAM7, promote collagen degradation, to regulate the survival, growth, migration, and invasion of cancer cells [49,50]. As collagen is one of the components of the cervix, it is suggested that this gene also acts in this organ, causing modifications. During cervical dilatation, collagen bundles are separated by increased water perfusion in the cervical extracellular membrane and collagen degradation by matrix metalloproteinases [51]. Kershaw et al. (2007) e Rodríguez-Piñón et al. (2015) [52,53] suggest that the breakdown of collagen fibers occurs from the late luteal phase until ovulation, followed by enzymatic collagen degradation. This gene is also highlighted among the 100 genes most expressed in the luteal phase of sheep (Table S4).

In our analysis, Hoxb genes were found in both cattle and sheep and their regulation may be involved in the PI3K/Akt pathway. Hox genes are considered members of the homeotic transcription factor family, they are the ones that will specify and direct the formation of tissues in the embryo [9]. Hox genes encode a class of transcription factors that play an important role in patterning axial development in vertebrates [54,55]. Hoxb2 appears to be specifically involved in motor neuron development [56]. Hoxb9 controls the specification of thoracic skeletal elements and mammary gland development and the regulation of various human cancers [57–60]. Hoxb8, it has already been reported that its overexpression in hematopoietic progenitor cells, in the presence of high concentrations of IL-3, allows the generation of growth factor-dependent myeloid cell lines capable of self-renewal [61–63]. De las Heras-Saldana et al. (2019) [64] provided evidence that the differential expression of some Hox genes, such as Hoxb2, Hoxb4, Hoxb9, may be involved in muscle differentiation, where these genes seem to modulate the muscle fate of satellite cells during myogenesis. But overall, the role of these Hox proteins in the cervical epithelium is still unclear [65]. These Hoxb genes appear to be involved in several biological processes, such as the formation of organs and muscles [66]. How the cervix undergoes structural, chemical, and biological differences throughout the estrous cycle [1], these Hox genes can modulate some mechanisms of cervical tissue transformation.

The PI3K/Akt signaling pathway was found in both cattle and sheep and could be regulated differently in both species (Figures 1 and 2). PI3K/Akt is involved in different signaling processes, ranging from apoptosis to cellular metabolism processes [67]. The PI3K/Akt pathway appears prominently in both species and may be involved in signal transduction mechanisms in different systems and organs. For communication between cells to occur, signaling pathways are necessary to command and mediate various functions, whether migration, differentiation, proliferation, or death [67]. PI3K/Akt plays important roles in control and balance in multicellular organisms [67]. Some studies have already documented the PI3K/Akt pathway signaling cascade and its involvement in the regulation of cell survival and apoptotic inhibition [68], as well as acting in the uptake of glucose and consequent involvement in the metabolism [69,70]. The PI3K/Akt pathway has been reported as an important survival pathway of eukaryotic cells, with the Akt serine/threonine kinase considered the key signaling point [71]. Furthermore, activated Akt directs an anti-apoptotic signal, thus protecting even cancer cells [72]. Its involvement is also related to the regulation of the functional activity of several proteins, which triggers different biological responses [73–76]. Thus, phosphorylation or dephosphorylation can control or regulate specific biological processes, catalyzing enzymes, regulating ion channels, transcription factors, intracellular protein localization, cytoskeleton regulation, and receptor activity [77].

Some genes found in our study have already been reported to be involved in the PI3K/Akt pathway (TDGF1, Hoxb2, DRD2) [37,56,78]. However, we showed that PI3K/Akt may be modulating most of the genes presented, as it appears in the analysis as one of the most expressed pathways in both sheep and cattle. The Hoxb genes are also modulating or are modulated by the PI3K/Akt pathway in both species (Figures 5 and 6). As this pathway is related to several biological processes, it may be an interesting pathway in the regulation of cervical relaxation mechanisms. Its stimulation or blockage can trigger cervical changes, as there is a relationship with several genes that supposedly participate in the process of dilation of the cervix.

The key gene most expressed in the follicular phase in cattle was MUC1 (Table S1). This gene has a fundamental role in the cervical mechanism, as there is an interaction between cervical mucus and collagen cells, as this mucus contains specific glycoproteins and enzymes that can act directly or indirectly on collagen [19]. In addition, this gene encodes a membrane-bound protein that is a member of the mucin family, which plays an essential role in the formation of protective mucous barriers on epithelial surfaces [79]. Cervical mucus can vary in its properties according to the estrous phase, attributing to the physiological need for sperm transport, acting as an antimicrobial agent, and also being a natural barrier in the luteal phase of the cycle and during pregnancy [80]. Mucin genes are already reported to show a strong correlation with increased estradiol levels [81] and that high cervical MUC1 expression during estrogen dominance enhances its known role as an antimicrobial barrier [82,83] especially when the cervix is open.

The hub gene most expressed in the follicular phase of sheep was ESR1, while in the luteal phase, it was ESR2 (Table S2). The presence of ESR1 is the main determinant of the regulation of the OXTR gene in the endometrial epithelium [84]. These are involved in muscle relaxation and contraction [85]. ESR2 has been reported in all uterine and uteroplacental tissue compartments, with constant expression throughout early pregnancy [86]. These receptors play a fundamental role in dilatation, as estradiol is fully involved in the cervical relaxation mechanism, as it indirectly induces smooth muscle relaxation and extracellular matrix remodeling [51]. In addition, the increase in estrogens during the follicular phase the of estrus cycle ewes is associated with natural cervical relaxation in estrus [1].

More expressed in the follicular phase in cattle (Table S3), BPIFA2A is a cationic protein stored in leukocytes with potent anti-endotoxin activity and highly selective bactericidal effects on Gram-negative bacteria [87,88]. This protein is considered bactericidal [89], and its role may be related to antimicrobial activity in the cervical canal. In the luteal phase, AGT (Table S3) is a participant in the renin-angiotensin system (RAS) and is well known for its role in regulating blood pressure and body fluid homeostasis [90]. Its role seems to be involved in the vasoconstrictor effect, apoptosis, angiogenesis and cell proliferation in several cell types [90]. In sheep, AGT encodes an angiotensin II product, described as a vasoconstrictor and regulator of fetoplacental angiogenesis in the placenta [91]. In the cervix, its role has not yet been elucidated. But it is already reported that there is an effect of angiotensin on the production of estradiol, however this mechanism is still not fully understood [92,93]. According to Giani et al. (2007) e Sampaio et al. (2007) [94,95] angiotensin may play a role in estradiol production through PI3K/Akt signaling.

Our analyzes indicate pathways and genes that may act directly on cervical relaxation. The PI3K/Akt pathway and the Hoxb genes are highlighted and can modulate and be modulated in mechanisms that involve alterations in the cervix. A more detailed study with blocking or stimulating this pathway and these genes in the cervix should be performed to assess cervical dilation during the luteal phase.

## 4. Conclusion

Differently expressed genes and divergent biological processes indicate differences related to cervical relaxation in cattle and sheep. PI3K/Akt appears to be an important pathway in the cervical relaxation process. *In vivo* studies of blockade or stimulation of this pathway should be performed to evaluate cervical relaxation in the estrous phase of sheep.

## 5. Materials and Methods

### 5.1 Bioinformatics for obtaining and processing data

The bovine cervix transcriptomic data in the follicular and luteal phase were provided by Pluta et al. (2012) [81] and sheep by Abril-Parreño [96], with the generated and/or analyzed datasets available in the NCBI Gene Expression Omnibus (https://www.ncbi.nlm.nih.gov/geo/) under accession number GSE38225 and GSE179486, respectively.

As a criterion for data selection, only samples of studies that contained information related to the sequencing platform used were adopted, giving preference to studies that made them available in “raw” form, containing physiological information regarding the samples and putting an end to the quality of the data. available data.

The files were downloaded in SRA format directly from GEO to the cluster of the Animal Improvement and Biotechnology Group at FZEA-USP. All data had the sequencing quality verified by the FASTQC software, followed by the removal of reads according to the data quality (“trimming”) with the TRIM GALORE software, both software from the Brabahan Institute. (https://www.bioinformatics.babraham.ac.uk/projects/index.html).

After selecting the reads by quality, we checked the quality with the R fastqcr package [97]. The samples that passed the quality test were then aligned with the reference genome of the species available at ENSEMBL (https://www.ensembl.org/info/data/ftp/index.html), with the RSUBREAD software [98], using the software’s default parameters suitable for each type of sample library. Alignment quality was then verified and a final report was generated using MULTIQC software [99].

### 5.2 Data Analysis, identification of gene signatures, and differences in expression

The analysis of gene expression difference was performed with the DESEQ2 package [100] of the R software [101] and exploratory data analysis was performed through principal component analysis (PCA) to the contrasts between the physiological situations of the cervix and between species.

For a gene to be considered differentially expressed, an adjusted p-value lower than 0.1 (padj <0.1) was adopted, by the Benjamini-Hochberg (“BH”) method, and an absolute value of “log2 fold change” greater than 2. For the representation of the gene expression values, the variance normalization transformation was used (function “varianceStabilizingTransformation” of the DESEQ2 package) which also served as an input for the data for the analysis of gene co-expression.

Gene co-expression analyzes were used to search for transcriptional profiles, “gene signatures”, in samples at different physiological phases of the cervix and species. In this phase, we used the CeTF package [102].

After gene selection, gene ontology analysis and pathway enrichment were performed with the ClusterProfiler package [103] of the R software. At first, we investigated the biological and molecular functions and the cellular components involved, in addition, we investigated the cured pathways in KEGG (“Kyoto Encyclopedia of Genes and Genomes”) and REACTOME. To visualize the results, networks of gene interaction and pathways were built using ClusterProfile.

## Author Contributions

The work was planned by J.D.G., R.P.N., M.E.F.O and performed by J.D.G and R.P.N., J.D.G wrote the manuscript and R.P.N., M.E.F.O., M.E.F.O, J.B.S.F and F.V.M resources, funding and critical revision of the manuscript. All authors read and agreed with the submitted version of the manuscript.

## Institutional Review Board Statement

Ethical review and approval were waived for this study, as data already published in journals were used.

## Data Availability Statement

Data Availability Statement: The data that support the findings of this study are available from the corresponding author upon reasonable request.

## Acknowledgments

The authors would like to thank the Coordination for the Improvement of Higher Education Personnel (CAPES) for funding the work. The team at the Molecular Morphophysiology and Development Laboratory of the Faculty of Food Engineering-FZEA at the University of Sao Paulo for their support.

