## Supplementary Materials for "An exploratory data analysis from ovine and bovine RNA-seq identifies pathways and key genes related to cervical dilatation"

**Table S1:** Hub genes in the bovine cervix in the follicular and luteal phases.

| Gene | ID | **Log Fold Change** | **Dif Means** | gini_1 | gini_2 | Mean1 | Mean2 |
|---|---|---|---|---|---|---|---|
| ENSBTAG00000046878 | ENSBTAG00000046878 | 0.323090905795746 | 0.0974687086131434 | 0.235706990262491 | 0.150256817634492 | 2.12500164212631 | 2.02753293351317 |
| HMG20A | ENSBTAG00000020441 | -0.097811620375919 | -35.6041149551268 | 0.150758398078968 | 0.055365825614059 | 223.899762583899 | 259.503877539026 |
| ZNF419 | ENSBTAG00000017613 | -0.554027692420169 | -4.47580907197521 | 0.194980575222539 | 0.0806871481349731 | 7.82256686356427 | 12.2983759355395 |
| KDM2A | ENSBTAG00000044032 | -0.278163633888021 | -28.8356191176694 | 0.0676107738152934 | 0.0164675188247649 | 70.3885376362167 | 99.224156753886 |
| MLF1 | ENSBTAG00000004126 | -0.325266641627905 | -0.232716529183624 | 0.293391867988911 | 0.125574877185134 | 0.829843253362801 | 1.06255978254643 |
| EAF1 | ENSBTAG00000019202 | 0.153662877986998 | -0.68872834124987 | 0.0755208628590565 | 0.0598917016902881 | 17.4318724379855 | 18.1206007792354 |
| ATXN1L | ENSBTAG00000046255 | -0.386820917363827 | -58.7855538815191 | 0.129073736081567 | 0.0708260078213465 | 105.67588459104 | 164.461438472559 |
| PLCB1 | ENSBTAG00000008338 | 0.0263776314191702 | 0.25523379759435 | 0.3722674457735 | 0.225286511119357 | 11.4034538041402 | 11.1482200065458 |
| MTA3 | ENSBTAG00000004073 | -0.737863953960057 | -1.02136351434001 | 0.166499515235935 | 0.0333845024301732 | 1.30144418349839 | 2.3228076978384 |
| CHD4 | ENSBTAG00000014734 | -0.130782859183929 | -6.78320930715659 | 0.0832530342936934 | 0.0334759724568583 | 24.6016587547516 | 31.3848680619082 |
| SERTAD3 | ENSBTAG00000010502 | 0.137178762257812 | -3.30498776424935 | 0.119601097591744 | 0.0620024387453041 | 41.5836528288874 | 44.8886405931367 |
| PAXIP1 | ENSBTAG00000017505 | -0.402946136882788 | -2.80396781033697 | 0.0995477836732697 | 0.0366688036738553 | 4.9062910235212 | 7.71025883385816 |
| CNOT11 | ENSBTAG00000008267 | -0.241937023687223 | -3.47670020759057 | 0.061492036316046 | 0.0804970145539508 | 8.41602899555025 | 11.8927292031408 |
| HDAC3 | ENSBTAG00000017360 | 0.224215468342189 | -0.0142271687011091 | 0.0659028887210294 | 0.0560509519930072 | 4.01614752448022 | 4.03037469318133 |
| PPP1R13L | ENSBTAG00000020689 | -0.0700090234373506 | -4.54311978602093 | 0.148848115499664 | 0.0762984316276214 | 17.9724270886137 | 22.5155468746347 |
| MED24 | ENSBTAG00000021468 | -0.286137172499933 | -11.3832112785057 | 0.174332595167555 | 0.0953427522916272 | 25.4681736958781 | 36.8513849743838 |
| MAPK13 | ENSBTAG00000010007 | 0.660204589458536 | 4.16958148725283 | 0.241961568190942 | 0.0805168388596126 | 15.887714391138 | 11.7181329038852 |
| DNM2 | ENSBTAG00000013362 | -0.284789014273538 | -56.6815812569633 | 0.117254565300117 | 0.0497941096933369 | 132.8287126338 | 189.510293890764 |
| POLR2G | ENSBTAG00000009483 | -0.187446185805971 | -25.7066726820744 | 0.0910016855977341 | 0.0702135471573257 | 78.314586446408 | 104.021259128482 |
| HDGF | ENSBTAG00000039793 | 0.320506261786834 | 5.48429561132043 | 0.182160554072257 | 0.0155602034016393 | 104.905183948395 | 99.4208883370741 |
| NCOA7 | ENSBTAG00000046503 | 0.15382574209718 | 0.173062698618675 | 0.271179627708595 | 0.0973093319484125 | 2.000303035604 | 1.82724033698532 |
| POLR1A | ENSBTAG00000006481 | -0.251278069173389 | -2.41805053225654 | 0.154504542773181 | 0.0313159547267263 | 5.85852213794489 | 8.27657267020144 |
| PCBD1 | ENSBTAG00000011866 | -1.00054718470556 | -55.2553850397195 | 0.0998312648474659 | 0.162383011750733 | 40.7776929514488 | 96.0330779911683 |
| FHL2 | ENSBTAG00000001086 | 0.485372105894402 | 1.94472104627881 | 0.136606376657373 | 0.086493415258272 | 11.2951894937647 | 9.35046844748589 |
| PARP1 | ENSBTAG00000000837 | -0.363783527108227 | -15.4769191237788 | 0.0398020461504979 | 0.0849686302412875 | 32.177613854876 | 47.6545329786548 |
| CTNND1 | ENSBTAG00000002411 | -0.248202512839939 | -32.9118857998668 | 0.0413229709870337 | 0.0641898744322226 | 87.4259484940269 | 120.337834293894 |
| HMGB2 | ENSBTAG00000015101 | 0.0951912890051601 | 0.00753577861681576 | 0.259541951499165 | 0.0716464616677119 | 1.7657280372761 | 1.75819225865928 |
| GTF2IRD1 | ENSBTAG00000006212 | 0.151755402219555 | 0.0407206459938303 | 0.105740675175251 | 0.0323560133039303 | 6.29552455959678 | 6.25480391360295 |
| NCOR1 | ENSBTAG00000013271 | -0.244658789238636 | -27.0169447226311 | 0.021559098048567 | 0.0720353525779446 | 72.102492583073 | 99.1194373057041 |
| PSMD9 | ENSBTAG00000004179 | -0.0370806282632892 | -18.7471768141625 | 0.110027436113434 | 0.07130251572394 | 82.0294603913207 | 100.776637205483 |

| | | | | | | | |
|---|---|---|---|---|---|---|---|
| AJUBA | ENSBTAG00000012724 | -0.529155495055643 | -1.61279835853387 | 0.208629847040172 | 0.0818242218433426 | 2.18200704736216 | 3.79480540589603 |
| CBX8 | ENSBTAG00000009107 | -0.210447734296982 | -1.46510266876569 | 0.172042185294599 | 0.079220029624588 | 3.70213479634887 | 5.16723746511456 |
| UXT | ENSBTAG00000015820 | 0.827741369211141 | 0.548291658019708 | 0.0934388038067035 | 0.0479168121332011 | 1.54252077195433 | 0.994229113934617 |
| DNMT3A | ENSBTAG00000021143 | -0.434134133243991 | -8.14509367130768 | 0.0567895353290105 | 0.0780697518952753 | 14.0972897830828 | 22.2423834543905 |
| MUC1 | ENSBTAG00000017104 | 1.5759491672159 | 42.8763687088901 | 0.223296975494568 | 0.121048657094685 | 69.9981017066941 | 27.1217329978039 |
| CASZ1 | ENSBTAG00000019818 | -0.476760960310648 | -13.6617530505828 | 0.179112085831366 | 0.0539784354556604 | 21.6177863357526 | 35.2795393863354 |
| AKAP8L | ENSBTAG00000010439 | -0.830693748870828 | -15.8819429664438 | 0.0754461648341227 | 0.0822111796622625 | 14.4085390060616 | 30.2904819725054 |
| TERF2IP | ENSBTAG00000015686 | 0.318015273128463 | 24.8849738754909 | 0.155105751275222 | 0.0616124817431729 | 192.18388228105 | 167.298908405559 |
| ERBB2 | ENSBTAG00000021798 | -0.292809880067697 | -10.6401816776785 | 0.175583223517799 | 0.0542535757904137 | 24.4191302538498 | 35.0593119315283 |
| HOXB5 | ENSBTAG00000045835 | 0.00864829046434397 | -1.04952052333382 | 0.225358852349498 | 0.067074862889544 | 6.37975612024244 | 7.42927664357627 |
| MRTFB | ENSBTAG00000008728 | -0.624873241342757 | -190.379788029047 | 0.0992043214792029 | 0.0742666803137045 | 265.581270076209 | 455.961058105256 |
| PRR13 | ENSBTAG00000026916 | 0.415231450983471 | 3.46838948028413 | 0.0976592569056149 | 0.0665391345487741 | 24.3860100653748 | 20.9176205850907 |
| TAF5L | ENSBTAG00000034225 | 0.00685040867780777 | -1.56754730946581 | 0.0698815481371179 | 0.0262408554417917 | 8.90813267844142 | 10.4756799879072 |
| LBH | ENSBTAG00000026111 | 1.62876889195243 | 68.3472801677114 | 0.195756522893946 | 0.206335254191895 | 111.832884919684 | 43.4856047519724 |
| KDM5B | ENSBTAG00000006175 | -0.0384307613041076 | -93.2508657006142 | 0.0526802723220755 | 0.0472437077893247 | 522.471897421563 | 615.722763122177 |
| HIPK2 | ENSBTAG00000017860 | -0.517739699942934 | -0.687623003232602 | 0.200709874829096 | 0.154832128115389 | 0.961722414598122 | 1.64934541783072 |
| RBPMS | ENSBTAG00000033727 | -0.0821872535967167 | -3.3341276480752 | 0.181214002198717 | 0.0477743446953832 | 17.2679523423757 | 20.6020799904509 |
| SUFU | ENSBTAG00000021068 | -0.430880083902824 | -48.333046361498 | 0.152733752122834 | 0.0565977788653143 | 82.2086505893704 | 130.541696950868 |
| DDIT3 | ENSBTAG00000031544 | 0.16505738512273 | -0.0151195362049265 | 0.0897820366710559 | 0.0710138370339491 | 0.570553272993806 | 0.585672809198732 |
| SIN3A | ENSBTAG00000009985 | -0.330965972069626 | -199.903782856967 | 0.0237860262980164 | 0.0423392114721217 | 438.465354755268 | 638.369137612235 |
| TTF2 | ENSBTAG00000015392 | 0.0335898096923506 | -0.184857914906793 | 0.0783349867149877 | 0.0942793689211042 | 1.45788406468644 | 1.64274197959323 |
| GLI3 | ENSBTAG00000010671 | -0.489594438274782 | -1.10526185431398 | 0.112982120825763 | 0.206483303510563 | 2.16201057768916 | 3.26727243200314 |
| WWTR1 | ENSBTAG00000007814 | 0.336308362701791 | 2.32037189128911 | 0.15467211358249 | 0.130324703370644 | 15.9233777818065 | 13.6030058905174 |
| MED27 | ENSBTAG00000000382 | -0.219793153551425 | -1.2610014618665 | 0.0864156924314682 | 0.0700801254637184 | 3.29738598755132 | 4.55838744941782 |
| NIF3L1 | ENSBTAG00000018282 | 0.0452452934566299 | -0.253769631446751 | 0.097878942781277 | 0.0746797978984681 | 1.97507604923806 | 2.22884568068481 |
| ZMYND8 | ENSBTAG00000013114 | 0.0407214186397831 | -4.79767537392712 | 0.103737274353588 | 0.0880773620047245 | 37.4699268041671 | 42.2676021780943 |
| AKAP8 | ENSBTAG00000001807 | -0.739349226986793 | -27.0594230934803 | 0.0360170729294317 | 0.116894580925637 | 28.9289675219679 | 55.9883906154482 |
| PIR | ENSBTAG00000009477 | -0.769558551092231 | -1.97221922120661 | 0.146186261809711 | 0.274238516276698 | 1.8700384879575 | 3.84225770916411 |
| CBX5 | ENSBTAG00000006246 | -0.200690663869542 | -1.26864230591765 | 0.234335433665849 | 0.275105125962369 | 3.13939720403573 | 4.40803950995339 |
| CHCHD2 | ENSBTAG00000040295 | 0.198797918675193 | 0.111341607413301 | 0.0273742790011501 | 0.0231861619172467 | 16.7999379301479 | 16.6885963227346 |
| EYA3 | ENSBTAG00000043989 | -0.781036767689086 | -4.17243304911453 | 0.108507282862601 | 0.188914797769056 | 4.23290512769502 | 8.40533817680956 |
| ERCC3 | ENSBTAG00000020777 | -0.301128827259608 | -2.23947701494021 | 0.0584551881090467 | 0.0616981470877299 | 5.37139029961986 | 7.61086731456008 |
| TTF1 | ENSBTAG00000018710 | -0.640128183027747 | -2.85342180007701 | 0.165523813941282 | 0.143836730900824 | 3.51149834479708 | 6.36492014487409 |
| CTBP2 | ENSBTAG00000003397 | -0.302457620831959 | -24.7988961520473 | 0.0257536836489903 | 0.118241347030093 | 56.7611236001937 | 81.560019752241 |
| ELOA | ENSBTAG00000026585 | -0.190045863234437 | -1.17622664833641 | 0.073625795492223 | 0.127380982139692 | 3.43279096938085 | 4.60901761771726 |

| | | | | | | | |
|---|---|---|---|---|---|---|---|
| NSD3 | ENSBTAG00000001529 | -0.40020208685063 | -2.72480938395963 | 0.120820044907004 | 0.191095218161158 | 5.12458954288114 | 7.84939892684077 |
| SETDB1 | ENSBTAG00000000098 | -0.137270406082548 | -2.54274015267468 | 0.0630142898147256 | 0.0464996484970122 | 10.3896364685647 | 12.9323766212394 |
| XRCC6 | ENSBTAG00000006103 | -0.0622779066596389 | -3.09018777209705 | 0.0565803523281743 | 0.0696450581495576 | 15.0992560366452 | 18.1894438087423 |
| EYA2 | ENSBTAG00000013336 | -0.526791657200815 | -33.2314654534435 | 0.221314203650821 | 0.194058006897459 | 48.5464792331013 | 81.7779446865448 |
| KMT2C | ENSBTAG00000024199 | -0.454799322092206 | -4.78000998816025 | 0.138748417396087 | 0.167401441851236 | 7.82611348466135 | 12.6061234728216 |
| ADNP2 | ENSBTAG00000005916 | -0.116163759756263 | -10.3260474111518 | 0.0413737094312209 | 0.0824022806520881 | 41.0894510663885 | 51.4154984775403 |
| PHF8 | ENSBTAG00000013289 | -0.278043931329241 | -0.560352132539939 | 0.0819503481526139 | 0.0691964908899573 | 1.37728642021192 | 1.93763855275186 |
| CHD7 | ENSBTAG00000021841 | 0.138016988592552 | -1.11585011467081 | 0.0574049388056796 | 0.118062717809342 | 16.1083784498887 | 17.2242285645595 |
| PRMT2 | ENSBTAG00000005503 | 0.223498490763962 | 0.477582624873879 | 0.0661436292350077 | 0.165762403066505 | 10.9413196563231 | 10.4637370314492 |
| ANP32A | ENSBTAG00000012365 | -0.545482901740455 | -12.360971099161 | 0.049446597918387 | 0.0820583915000928 | 18.4424967002574 | 30.8034677994184 |
| TLE4 | ENSBTAG00000003532 | -0.372693879438917 | -3.30437445747314 | 0.134462867061302 | 0.1435660025412 | 7.82102573885292 | 11.1254001963261 |
| TAF9B | ENSBTAG00000000895 | 0.620916141230957 | 2.54431516417144 | 0.116160062199806 | 0.0617225979069945 | 9.38427087439089 | 6.83995571021944 |

**Table S2:** Hub genes in the sheep cervix in the follicular and luteal phases.

| Gene | ID | **Log Fold Change** | **Dif Means** | gini_1 | gini_2 | Mean1 | Mean2 |
|---|---|---|---|---|---|---|---|
| HOXB3 | ENSG00000120093 | -0.0444430789760876 | -0.314671750814491 | 0.151961424329116 | 0.0935109779778038 | 3.47504846716703 | 3.78972021798152 |
| ZNF34 | ENSG00000196378 | -0.675857205978606 | -8.6660830579716 | 0.162368238741957 | 0.127249655493742 | 12.6954084135128 | 21.3614914714844 |
| ESRRA | ENSG00000173153 | 0.359070099652815 | 1.62194048171043 | 0.153013490022434 | 0.104512468758234 | 9.55126023190595 | 7.92931975019552 |
| PREB | ENSG00000138073 | 0.0444018064928695 | -54.2517992481799 | 0.114569552788885 | 0.0607799165187727 | 1456.45163360534 | 1510.70343285352 |
| NFIL3 | ENSG00000165030 | -0.0169807492978161 | -0.249525902434344 | 0.152315853464092 | 0.16227380802779 | 2.93097132679661 | 3.18049722923096 |
| ERF | ENSG00000105722 | -0.0257949519021511 | -1.33169227757189 | 0.0732487953125763 | 0.0756843804643073 | 16.7055796454344 | 18.0372719230063 |
| ZFP69 | ENSG00000187815 | -0.0163665173166672 | -0.465628586900626 | 0.108476231114715 | 0.110361484286911 | 6.39583826781362 | 6.86146685471425 |
| NFATC4 | ENSG00000285485 | -0.0613435394215662 | -0.052688596466541 | 0.190533197525829 | 0.187300817469598 | 0.473455585261797 | 0.526144181728338 |
| HOXB2 | ENSG00000173917 | 0.12631164230233 | 0.0212668761551627 | 0.124500139406979 | 0.0872380906881615 | 0.858067660572667 | 0.836800784417504 |
| MEIS3 | ENSG00000105419 | -0.193128262018687 | -0.885761589655636 | 0.176077876350497 | 0.189835438101973 | 4.06396677084364 | 4.94972836049927 |
| EHF | ENSG00000135373 | 0.0551965703476608 | -1.67338329120975 | 0.196017368527114 | 0.215629831212362 | 66.8751223503233 | 68.548505641533 |
| CIZ1 | ENSG00000148337 | -0.131516997169159 | -14.9721076043833 | 0.0593695173976606 | 0.0521777490981299 | 93.0444150402619 | 108.016522644645 |
| ZNF750 | ENSG00000141579 | 0.0594469909348513 | -0.0322659848142932 | 0.220586172762167 | 0.257569746526407 | 1.54308108925488 | 1.57534707406917 |
| SPDEF | ENSG00000124664 | 0.0480488621999171 | -0.159760552038249 | 0.279871408616987 | 0.296337915564375 | 4.20658317236293 | 4.36634372440118 |
| SIX1 | ENSG00000126778 | -0.254342448908673 | -0.941013987856023 | 0.199799343567225 | 0.175946570997883 | 3.51375109242425 | 4.45476508028027 |
| MAF | ENSG00000178573 | 0.549186451698951 | 0.372364257677024 | 0.244690329596164 | 0.226006961680613 | 1.33099585461996 | 0.958631596942931 |
| NFX1 | ENSG00000086102 | -0.548530206406819 | -0.189164549938 | 0.0701254664301291 | 0.072639897843982 | 0.345328133745948 | 0.534492683683947 |
| ZNF276 | ENSG00000158805 | -0.121635166801796 | -2.87300000942937 | 0.0582528997995442 | 0.0606613116519825 | 18.5651606480843 | 21.4381606575137 |
| SMARCE1 | ENSG00000073584 | 0.196547198465193 | 0.193985480805279 | 0.0589504960384857 | 0.0466811287881742 | 2.67820622279822 | 2.48422074199294 |
| ZNF652 | ENSG00000198740 | -0.155346848428834 | -3.85356440022137 | 0.12897091404807 | 0.18332601432775 | 21.0685773436668 | 24.9221417438882 |
| GLIS3 | ENSG00000107249 | -0.749938080238254 | -0.0820176207717817 | 0.160327753176676 | 0.234603031391278 | 0.104947898379246 | 0.186965519151028 |
| ZNF710 | ENSG00000140548 | -0.275250488787159 | -0.449058234688207 | 0.120555352842691 | 0.0637561280594882 | 1.59255599435058 | 2.04161422903879 |
| NFKB2 | ENSG00000077150 | 0.0739894498970104 | -0.0609063251642361 | 0.154318775631769 | 0.16045039234085 | 13.8668550230431 | 13.9277613482074 |
| MYC | ENSG00000136997 | -0.220300417128693 | -207.15784306916 | 0.135040985351753 | 0.130502425908539 | 866.605574159048 | 1073.76341722821 |
| ESR1 | ENSG00000091831 | 1.01079315236001 | 60.3619813538215 | 0.173406092548588 | 0.188435638047441 | 128.045272456939 | 67.6832911031179 |
| PRDM2 | ENSG00000116731 | -0.442546886518727 | -40.3011041728139 | 0.115748330406608 | 0.228262116953901 | 91.2567397626774 | 131.557843935491 |
| STAT6 | ENSG00000166888 | 0.00992095296381899 | -0.780117575573508 | 0.0835705175230118 | 0.0855312731787687 | 15.5401156278796 | 16.3202332034531 |
| PKNOX2 | ENSG00000165495 | 0.340110364637196 | 0.537233345695989 | 0.186571725957367 | 0.269787541268671 | 3.35377204074548 | 2.81653869504949 |
| ESR2 | ENSG00000140009 | -1.10402057614656 | -4.06063822919675 | 0.449806144012118 | 0.49348544621593 | 3.19638564515389 | 7.25702387435064 |
| NFKB1 | ENSG00000109320 | 0.0149791993426645 | -44.1303882919558 | 0.110342446846259 | 0.0973525054412114 | 1005.12136278387 | 1049.25175107582 |
| TP53 | ENSG00000141510 | 0.199871045564648 | 0.350397278968463 | 0.107219555382859 | 0.108659714239451 | 4.66407760591478 | 4.31368032694632 |
| MEF2D | ENSG00000116604 | 0.209997237112109 | 0.292505529228725 | 0.0710573153580517 | 0.0850085171579094 | 3.46446678143662 | 3.1719612522079 |
| ETS1 | ENSG00000134954 | 0.018217625848141 | -0.0500412714152569 | 0.138495376636028 | 0.181634901701882 | 1.01989553583063 | 1.06993680724589 |

**Table S3:** The 100 most expressed genes in the follicular and luteal phase of the bovine cervix.

| Gene | ID | **Log Fold Change** | **Dif Means** | gini_1 | gini_2 | Mean1 | Mean2 |
|---|---|---|---|---|---|---|---|
| BPIFA2A | ENSBTAG00000009144 | 10.4876703575293 | 1.33314666767432 | 0.536551689835304 | 0.8 | 1.33379545013532 | 0.000648782460996904 |
| TMPRSS11BNL | ENSBTAG00000048377 | 9.13963845167615 | 44.6748103458901 | 0.492638564074233 | 0.416231380534227 | 44.7637138270306 | 0.0889034811405015 |
| ENSBTAG00000048276 | ENSBTAG00000048276 | 8.83451158367456 | 11.0798911087455 | 0.260716716149161 | 0.368647838134362 | 11.1076696230359 | 0.0277785142904404 |
| LOC112441508 | ENSBTAG00000031375 | 8.43389967371738 | 1.48923886546534 | 0.528562019526543 | 0.623714018536848 | 1.49442237481876 | 0.00518350935341578 |
| BPIFA2B | ENSBTAG00000019752 | 8.01169488659356 | 0.627058853372568 | 0.495585367076419 | NA | 0.627058853372568 | 0 |
| AGT | ENSBTAG00000012393 | -7.72323486386274 | -2.83968274534543 | 0.391639105233969 | 0.769482777513028 | 0.0107830276558119 | 2.85046577300124 |
| TMPRSS11D | ENSBTAG00000001925 | 7.2661091230698 | 5.64468362774846 | 0.535834114542948 | 0.401650565146232 | 5.68581144313726 | 0.0411278153887952 |
| CAMK2B | ENSBTAG00000012653 | -7.19841629076593 | -36.6400263143898 | 0.481605736403381 | 0.775229010285204 | 0.209178053159811 | 36.8492043675496 |
| TMIE | ENSBTAG00000052515 | -7.14496785402895 | -32.2320531948505 | 0.746704300909962 | 0.271334587578201 | 0.166251472053501 | 32.398304666904 |
| NCCRP1 | ENSBTAG00000014296 | 7.11259157979939 | 1.28225291262756 | 0.486041477498022 | 0.615406405211467 | 1.29406602372885 | 0.0118131111012905 |
| OLFM4 | ENSBTAG00000022779 | 6.98174767844643 | 13.837555827811 | 0.308109841450442 | 0.689536041445065 | 13.9525652777417 | 0.11500944993065 |
| TDGF1 | ENSBTAG00000021119 | -6.80310364038778 | -201.057849100285 | 0.310916583251406 | 0.631884126277704 | 1.65549732489853 | 202.713346425183 |
| ENSBTAG00000011470 | ENSBTAG00000011470 | -6.77458040455884 | -10.1650175378062 | 0.446871761062352 | 0.140478886939971 | 0.0729406735126094 | 10.2379582113188 |
| PADI1 | ENSBTAG00000002138 | 6.65159901658234 | 25.1419413304667 | 0.412782797072968 | 0.355129926047248 | 25.4312967007317 | 0.289355370265031 |
| KRT13 | ENSBTAG00000050581 | 6.60455651689265 | 48.8673317699611 | 0.47304553549609 | 0.518094262785813 | 49.4588326497406 | 0.591500879779518 |
| SCG2 | ENSBTAG00000021588 | 6.46281630849971 | 0.828607699462793 | 0.713710958282025 | 0.8 | 0.834794095323741 | 0.00618639586094831 |
| LRP2 | ENSBTAG00000004555 | -6.4320396209309 | -9.73441966584519 | 0.336120178665147 | 0.563394511740939 | 0.0974085128359711 | 9.83182817868116 |
| ENSBTAG00000053748 | ENSBTAG00000053748 | 6.30191233312157 | 53.7587482952646 | 0.556960222428914 | 0.39367890594909 | 54.5359006047368 | 0.777152309472178 |
| KRT4 | ENSBTAG00000012034 | 6.30006056405355 | 2.8479905970231 | 0.557329403416542 | 0.39367890594909 | 2.88921883625239 | 0.0412282392292932 |
| GAL | ENSBTAG00000009393 | 6.25065974277691 | 332.298429497016 | 0.250359346249565 | 0.442329997859117 | 337.161000271159 | 4.86257077414229 |
| SLC30A8 | ENSBTAG00000052098 | -6.20793360598155 | -11.1683475480057 | 0.576497311585583 | 0.616842353894826 | 0.116853247037114 | 11.2852007950428 |
| MGC138914 | ENSBTAG00000014328 | -6.20252448589179 | -131.701416485042 | 0.511629942801625 | 0.290781077446007 | 1.53555954194665 | 133.236976026989 |
| UPK2 | ENSBTAG00000012750 | 6.17135102341042 | 0.509952115161376 | 0.590019242974515 | 0.635711877517133 | 0.51822726966561 | 0.00827515450423429 |
| LOC522479 | ENSBTAG00000050427 | 6.13045929224964 | 154.713158413611 | 0.434115962813147 | 0.319281627009887 | 156.990223838749 | 2.27706542513746 |
| ENSBTAG00000055199 | ENSBTAG00000055199 | -6.11874699133316 | -1.21966498637771 | 0.671609886916306 | 0.780052974541294 | 0.0130108581114812 | 1.23267584448919 |
| IHH | ENSBTAG00000008452 | -6.06063859427148 | -6.41313863037376 | 0.50762878188206 | 0.790167344499239 | 0.0823521556816896 | 6.49549078605545 |
| F10 | ENSBTAG00000016385 | -5.97778670464108 | -1.82341483696047 | 0.722329456151623 | 0.631340237387627 | 0.0227455913127713 | 1.84616042827325 |
| REG4 | ENSBTAG00000032193 | 5.97254369580806 | 2.76426651309394 | 0.457878650885152 | 0.494784534101104 | 2.81112852797451 | 0.046862014880569 |
| LOC112446672 | ENSBTAG00000052798 | 5.97028153990243 | 0.422175917456848 | 0.568501932138137 | NA | 0.422175917456848 | 0 |
| SPINK1 | ENSBTAG00000015558 | 5.97021056305912 | 2.92933404999975 | 0.427337973985977 | 0.23488985350991 | 2.98042556807162 | 0.0510915180718691 |
| SLURP1 | ENSBTAG00000016209 | 5.96628107842237 | 2.13890518318835 | 0.462899263017627 | 0.468517291928679 | 2.17925251219645 | 0.0403473290080999 |
| MSMB | ENSBTAG00000011660 | 5.9172246051464 | 11.559692840123 | 0.57438559357943 | 0.420535691388361 | 11.7680586379666 | 0.208365797843637 |
| MMP3 | ENSBTAG00000037768 | 5.90784242672446 | 0.300619796967037 | 0.575586118756744 | 0.687145987100115 | 0.305441167173454 | 0.0048213702064169 |
| LOC101906048 | ENSBTAG00000046283 | -5.81406656406624 | -112.22982739435 | 0.532964803569153 | 0.640889438458854 | 1.6756578031257 | 113.905485197476 |
| TAAR1 | ENSBTAG00000037718 | 5.69125738448153 | 5.30173948963706 | 0.350309420495675 | 0.338114887682562 | 5.41421185107781 | 0.112472361440749 |
| BPIFA2C | ENSBTAG00000031376 | 5.66134798079537 | 6.93276385461759 | 0.439228909687748 | 0.198886695611993 | 7.08485654903631 | 0.152092694418717 |

| | | | | | | | |
|---|---|---|---|---|---|---|---|
| EXD1 | ENSBTAG00000008363 | 5.64090424776476 | 3.62173406094445 | 0.442225725513415 | 0.223980633222556 | 3.70258022965377 | 0.0808461687093241 |
| KCNF1 | ENSBTAG00000021280 | -5.56880608077551 | -0.733886478768648 | 0.588670714403363 | 0.3056205133675 | 0.0120916607764349 | 0.745978139545083 |
| TMPRSS3 | ENSBTAG00000048816 | 5.50926031435089 | 233.547919228969 | 0.296184512776969 | 0.523349310076629 | 238.835256077509 | 5.28733684853941 |
| ENSBTAG00000048816 | ENSBTAG00000020512 | 5.49427315123949 | 4.03539223846517 | 0.438178638298546 | 0.604198292230674 | 4.13164007802351 | 0.0962478395583349 |
| GJB1 | ENSBTAG00000008161 | -5.46737867462709 | -7.7516972071287 | 0.513730635930614 | 0.573761494647608 | 0.147320164154682 | 7.89901737128338 |
| CLCA1 | ENSBTAG00000052224 | 5.46220414840613 | 216.918610691352 | 0.348520323717683 | 0.220911275830118 | 222.367665187114 | 5.44905449576143 |
| ENSBTAG00000052224 | ENSBTAG00000046768 | -5.44645341710979 | -0.141385399020172 | 0.732679109278405 | 0.592853116464926 | 0.00158103991329466 | 0.142966438933466 |
| IGFBP1 | ENSBTAG00000031532 | 5.41407486190766 | 0.232438603880333 | 0.398708784245792 | 0.687781598208397 | 0.237886522437604 | 0.00544791855727039 |
| ENSBTAG00000031532 | ENSBTAG00000001392 | -5.30610654888893 | -9.21065752901083 | 0.392048787077624 | 0.729561457520693 | 0.198741578938214 | 9.40939910794904 |
| RDH16 | ENSBTAG00000014683 | -5.30278893747946 | -0.211260754246596 | NA | 0.22153142686123 | 0 | 0.211260754246596 |
| APOBEC1 | ENSBTAG00000038835 | 5.14904004851144 | 1.7300673491015 | 0.529195805451826 | 0.620588665099255 | 1.78304759578754 | 0.0529802466860465 |
| LOC509961 | ENSBTAG00000004588 | 5.125849712773 | 0.51405456520375 | 0.670915502609821 | 0.413939241131895 | 0.533021604361805 | 0.018967039158055 |
| KCNN4 | ENSBTAG00000022246 | 5.11845131628365 | 9.31715085297592 | 0.271822638609792 | 0.122277235259634 | 9.63955600286589 | 0.322405149889969 |
| C29H11orf86 | ENSBTAG00000018703 | 5.11490131443356 | 0.901634401094783 | 0.466235913936583 | NA | 0.901634401094783 | 0 |
| OSTN | ENSBTAG00000039237 | 5.10472743361731 | 2.13381832461088 | 0.493917195234149 | 0.619924271681624 | 2.1905700177357 | 0.0567516931248251 |
| ENSBTAG00000039237 | ENSBTAG00000050026 | -5.0899259170152 | -0.466797520303263 | 0.857142857142857 | 0.748126166762856 | 0.00991140105104993 | 0.476708921354313 |
| ENSBTAG00000050026 | ENSBTAG00000047155 | 5.08257740147134 | 0.10570344761818 | 0.366920338844249 | 0.8 | 0.108275364342786 | 0.00257191672460578 |
| C28H10orf71 | ENSBTAG00000012621 | 5.05666166554248 | 0.623191000853263 | 0.430556596569112 | 0.608384191973237 | 0.643170830812637 | 0.0199798299593742 |
| RTN4RL2 | ENSBTAG00000039967 | -5.02169556579284 | -4.52817025221992 | 0.372114164867171 | 0.191777165832691 | 0.112258073916326 | 4.64042832613624 |
| KRT78 | ENSBTAG00000021240 | 5.01230629889222 | 5.32986970531806 | 0.543476717946431 | 0.33561895102292 | 5.5235494764874 | 0.193679771169345 |
| DCSTAMP | ENSBTAG00000019636 | -4.99771198991172 | -1.75079853951409 | 0.595687858980708 | 0.370993004192074 | 0.0303553412939253 | 1.78115388080802 |
| SCARA5 | ENSBTAG00000039289 | -4.99333979615527 | -175.992664251122 | 0.34694681929318 | 0.376336987526026 | 5.40145840614963 | 181.394122657272 |
| LOC527068 | ENSBTAG00000053443 | 4.9914602241738 | 4.2131051169979 | 0.557702964942814 | 0.243677868243968 | 4.36668800060571 | 0.153582883607807 |
| RUM1 | ENSBTAG00000021280 | 4.98204833007017 | 9.74751598331487 | 0.150582097847591 | 0.262393338886956 | 10.0955034837624 | 0.347987500447485 |
| IVL | ENSBTAG00000017827 | 4.98047792619673 | 3.53860191933584 | 0.56752775113844 | 0.635711877517133 | 3.6725988335046 | 0.133996914168757 |
| ENSBTAG00000054600 | ENSBTAG00000054600 | 4.97711136426043 | 10.2692488853657 | 0.326277886234182 | 0.345770984835542 | 10.6364735005654 | 0.367224615199633 |
| CALML5 | ENSBTAG00000013854 | 4.91059617065609 | 0.46970608591465 | 0.244710652791082 | 0.684360546693485 | 0.488532537211412 | 0.0188264512967618 |
| LOC538679 | ENSBTAG00000046482 | 4.90470651814782 | 0.397143436621251 | 0.263156523890617 | 0.615406405211467 | 0.415176626600994 | 0.0180331899797425 |
| PENK | ENSBTAG00000004924 | -4.82018704800901 | -0.501637147293282 | 0.575389941722831 | 0.623033246461788 | 0.0142549202220615 | 0.515892067515344 |
| TH | ENSBTAG00000026768 | 4.81970880567878 | 13.392416503268 | 0.761638079877615 | 0.190218988191608 | 14.0002827140149 | 0.607866210746869 |
| A4GNT | ENSBTAG00000001451 | 4.80713439336761 | 0.300594879286845 | 0.435646804527321 | 0.258528707326984 | 0.314254997453264 | 0.0136601181664187 |
| ENSBTAG00000048830 | ENSBTAG00000048830 | 4.80228490133332 | 0.0270053431419432 | 0.423799663506785 | NA | 0.0270053431419432 | 0 |
| BCAN | ENSBTAG00000015789 | 4.75558678325738 | 1.39685057289354 | 0.293854499177153 | 0.212551789148209 | 1.46213645276024 | 0.0652858798666992 |
| CCL26 | ENSBTAG00000052510 | 4.74406346202816 | 10.5467762364544 | 0.166608731000911 | 0.149745425037113 | 11.0033532317772 | 0.456576995322828 |
| AGR2 | ENSBTAG00000024406 | 4.74106289947718 | 4060.3690683971 | 0.188915372048269 | 0.211978516408433 | 4235.21089557 | 174.841827172901 |
| ASPG | ENSBTAG00000017194 | -4.68688328813046 | -1.51479249813694 | 0.417312971349325 | 0.77426485970263 | 0.0486697215946323 | 1.56346221973158 |
| RADIL | ENSBTAG00000021913 | -4.67776436525556 | -7.40588386577895 | 0.266722263387651 | 0.767587755940789 | 0.251688861121091 | 7.65757272690004 |
| TMEM229A | ENSBTAG00000049382 | 4.66450793972288 | 0.192363626559942 | 0.441941247442293 | NA | 0.192363626559942 | 0 |
| KRT6A | ENSBTAG00000039425 | 4.65937157668344 | 4.52392082311196 | 0.525127285425684 | 0.529879305124812 | 4.73351117031102 | 0.209590347199066 |

| | | | | | | | |
|---|---|---|---|---|---|---|---|
| GJB5 | ENSBTAG00000005717 | -4.61265825636491 | -5.08705466735846 | 0.514698079451125 | 0.700279354678321 | 0.186697989645475 | 5.27375265700394 |
| ADM2 | ENSBTAG00000054072 | 4.61120314060221 | 14.0931763721036 | 0.226673923658365 | 0.192053192614435 | 14.7995669229185 | 0.70639055081487 |
| MCOLN3 | ENSBTAG00000016982 | -4.59926761289637 | -1.53294796619728 | 0.614935535631122 | 0.439745674542315 | 0.0472139550344666 | 1.58016192123174 |
| ENSBTAG00000005324 | ENSBTAG00000005324 | 4.58816104654107 | 2.82899586567018 | 0.572272110442103 | NA | 2.82899586567018 | 0 |
| IGFBP3 | ENSBTAG00000003994 | 4.58517460230098 | 95.2488081081069 | 0.0994427776122484 | 0.294491073326218 | 99.7007290097979 | 4.45192090169103 |
| DCT | ENSBTAG00000002300 | 4.56942007303733 | 0.148815689838838 | 0.781024732486731 | 0.8 | 0.15533952547402 | 0.00652383563518186 |
| BSP3 | ENSBTAG00000003886 | -4.55507383233905 | -0.640788200719888 | NA | 0.358662173213077 | 0 | 0.640788200719888 |
| ENSBTAG00000048395 | ENSBTAG00000048395 | -4.55475871869254 | -1.49175928917298 | 0.259681812333872 | 0.413284074808977 | 0.0679444069205873 | 1.55970369609357 |
| CYP3A4 | ENSBTAG00000052665 | 4.54947861024019 | 16.1959899325145 | 0.505185337093724 | 0.234091152098849 | 17.0043917722522 | 0.8084018397377 |
| KLF17 | ENSBTAG00000047871 | -4.51816238250645 | -1.18224713276154 | 0.679395546313255 | 0.232158498444094 | 0.0567615033820296 | 1.23900863614357 |
| TNC | ENSBTAG00000000575 | 4.43688586273517 | 208.62727582452 | 0.377655272939868 | 0.387453477402939 | 219.345984954238 | 10.7187091297178 |
| MGAT4C | ENSBTAG00000011153 | -4.42830804103556 | -10.1430414632334 | 0.444765811814567 | 0.393854200026938 | 0.457699191000382 | 10.6007406542338 |
| KCNE1 | ENSBTAG00000001150 | 4.41825885492678 | 0.283951444740196 | 0.667496459654903 | 0.8 | 0.29826234934035 | 0.0143109046001541 |
| C28H10orf99 | ENSBTAG00000050197 | -4.40459288804529 | -2.46284578703824 | 0.595524385461041 | 0.196760828426254 | 0.0925943051593968 | 2.55544009219764 |
| SFN | ENSBTAG00000009223 | 4.38739338650954 | 14.219610237416 | 0.493299337159911 | 0.156621523347076 | 15.0037883414765 | 0.784178104060512 |
| PRODH | ENSBTAG00000047676 | -4.37823848956379 | -8.38833718911427 | 0.165357870002746 | 0.24670042616056 | 0.360378147127657 | 8.74871533624193 |
| S100A12 | ENSBTAG00000012638 | 4.37468878868362 | 10.7508742455153 | 0.529950363454291 | 0.367741910941125 | 11.309453729421 | 0.558579483905632 |
| ENSBTAG00000052585 | ENSBTAG00000052585 | -4.37085733781731 | -4.65499638608366 | 0.529564529737433 | 0.566681904472133 | 0.172518301137926 | 4.82751468722159 |
| SLC14A1 | ENSBTAG00000019870 | 4.34841217170969 | 3.27389980244996 | 0.719294681521202 | 0.43878817341671 | 3.4405727956693 | 0.166672993219341 |
| GJB4 | ENSBTAG00000005719 | -4.3460233148246 | -9.2846910699956 | 0.455467368287012 | 0.441146566258235 | 0.390694171159466 | 9.67538524115507 |
| ENSBTAG00000048794 | ENSBTAG00000048794 | 4.34073589208589 | 167.677422864312 | 0.41365717422201 | 0.41744372950316 | 177.361731400924 | 9.68430853661206 |
| LOC112441481 | ENSBTAG00000012540 | -4.33881125665703 | -0.529486086435191 | 0.709098966057574 | 0.329646698327027 | 0.048095017148613 | 0.577581103583804 |
| LOC507527 | ENSBTAG00000013507 | 4.33049803539077 | 0.00796028178064769 | 0.562107108912761 | NA | 0.00796028178064769 | 0 |
| ENSBTAG00000053707 | ENSBTAG00000053707 | 4.3267015493717 | 1.58412444509021 | 0.40865370111719 | 0.444099991082275 | 1.67820105617526 | 0.0940766110850541 |
| S100A8 | ENSBTAG00000012640 | 4.31913547505644 | 1.19861552784099 | 0.5233191170937 | 0.38928140268617 | 1.26725393728352 | 0.0686384094425284 |

**Table S4:** The 100 most expressed genes in the follicular and luteal phase of the sheep cervix.

| Gene | ID | **Log Fold Change** | **Dif Means** | gini_1 | gini_2 | Mean1 | Mean2 |
|---|---|---|---|---|---|---|---|
| ENSOARG00020011601 | ENSOARG00020011601 | 7.65678771992481 | 2.33754749161547 | 0.767658467531659 | 0.735541205336112 | 2.35002171949169 | 0.0124742278762233 |
| ENSOARG00020000537 | ENSOARG00020000537 | 7.32707456273094 | 0.0753685444755734 | 0.81927013027439 | 0.736979654019151 | 0.0758769005296702 | 0.000508356054096724 |
| KRTDAP | ENSG00000188508 | 7.25380690371756 | 0.0111377364888888 | 0.858832754411478 | 0.839786875344018 | 0.0111839536402679 | 4.62171513791126e-05 |
| DRD2 | ENSG00000149295 | 6.7507378119605 | 51.670365850602 | 0.463983997794401 | 0.853983800457149 | 52.1954422878483 | 0.52507643724623 |
| SLURP1 | ENSG00000126233 | 6.64203575141894 | 0.258738719182664 | 0.672064308043247 | 0.802490451645851 | 0.261554540361375 | 0.00281582117871167 |
| KRT4 | ENSG00000170477 | 6.64118502059344 | 1070.81788636707 | 0.825671836911915 | 0.924309206282594 | 1115.30013891217 | 44.482252545096 |
| PLA2G4E | ENSG00000188089 | 6.57073948117742 | 0.594890875845066 | 0.870768628493842 | 0.875252767091832 | 0.596887198985571 | 0.00199632314050495 |
| ENSOARG00020018856 | ENSOARG00020018856 | 6.16812848914848 | 0.00738487708862217 | 0.892911726650831 | 0.697396395694228 | 0.00746780094259318 | 8.29238539710077e-05 |
| MMP1 | ENSG00000196611 | 6.16576596182193 | 0.403520062244577 | 0.676630866949708 | 0.894876015890183 | 0.40947511233817 | 0.00595505009359361 |
| TECTB | ENSG00000119913 | 5.83895349961854 | 0.130225827049176 | 0.73503534592538 | 0.696393706901843 | 0.132679945325323 | 0.00245411827614712 |
| SCG2 | ENSG00000171951 | 5.65339614183538 | 0.867067709064849 | 0.540959058728606 | 0.831792221054691 | 0.885957372812962 | 0.0188896637481134 |
| KRT79 | ENSG00000185640 | 5.63144322349349 | 2.28717664250687 | 0.74623858964369 | 0.767969205304519 | 2.33717117174681 | 0.0499945292399396 |
| CSN3 | ENSG00000171209 | 5.62539017807886 | 1.12964207762822 | 0.899556216605685 | 0.916461904836629 | 1.13716401976393 | 0.0075219421357065 |
| IBSP | ENSG00000029559 | 5.60558222079176 | 0.233812769438452 | 0.666056914977453 | 0.93875017133709 | 0.23617825645322 | 0.00236548701476805 |
| ADAM7 | ENSG00000069206 | -5.58250234721926 | -3.24039388002412 | 0.576242541355225 | 0.647289715558404 | 0.0654185677861933 | 3.30581244781032 |
| ENSOARG00020019270 | ENSOARG00020019270 | 5.56729164751594 | 0.146829870523219 | 0.955010580434334 | 0.817380378632078 | 0.147397625031332 | 0.000567754508112833 |
| ENSOARG00020010220 | ENSOARG00020010220 | -5.50926952410315 | -19.9830095240443 | 0.8423808079539 | 0.873602332007091 | 0.42262162265611 | 20.4056311467004 |
| ENSOARG00020012069 | ENSOARG00020012069 | 5.42856128341953 | 1.39833297284088 | 0.729774021844396 | 0.580709863778545 | 1.43393308416918 | 0.0356001113282972 |
| ENSOARG00020024490 | ENSOARG00020024490 | 5.42054576262215 | 0.201074255363184 | 0.85907376486467 | 0.834514981307652 | 0.206514545665752 | 0.00544029030256842 |
| KRT85 | ENSG00000135443 | -5.36888720716065 | -0.679769137156458 | 0.886335091400448 | 0.804029299987675 | 0.0144309064747076 | 0.694200043631165 |
| TFF1 | ENSG00000160182 | 5.31535865509055 | 2.77619608234766 | 0.673581585724974 | 0.709475018445018 | 2.85111580527535 | 0.0749197229276895 |
| ENSOARG00020011654 | ENSOARG00020011654 | 5.3040154599145 | 18.2264964452496 | 0.770570686141199 | 0.689710973149857 | 18.7332947083822 | 0.506798263132556 |
| ENSOARG00020015596 | ENSOARG00020015596 | -5.195565115895 | -1.29049973712559 | 0.887666111980304 | 0.84821302845308 | 0.0342772063486563 | 1.32477694347424 |
| ENSOARG00020000114 | ENSOARG00020000114 | 5.17939370895714 | 0.430830278181781 | 0.897266673501732 | 0.783973408492965 | 0.440758771530191 | 0.00992849334840928 |
| ENSOARG00020000372 | ENSOARG00020000372 | 5.10688287182257 | 0.0723453713139255 | 0.767810780167757 | 0.53851386930478 | 0.074657214528892 | 0.0023118432149665 |
| ENSOARG00020024493 | ENSOARG00020024493 | -5.02464664754408 | -1.01468859896477 | 0.416676548651069 | 0.669929118968995 | 0.030593915782405 | 1.04528251474717 |
| ENSOARG00020013969 | ENSOARG00020013969 | -4.99117002843005 | -0.0198389644560448 | 0.820235129618376 | 0.695395389932465 | 0.000601968203748093 | 0.0204409326597929 |
| ENSOARG00020012116 | ENSOARG00020012116 | 4.98002467282859 | 0.0279318533758113 | 0.857970026929464 | 0.942061535861 | 0.0282288172221723 | 0.000296963846360968 |
| ENSOARG00020011773 | ENSOARG00020011773 | 4.92106084845893 | 3.62959702073834 | 0.705832452761686 | 0.552839104353695 | 3.76124504961823 | 0.131648028879884 |
| SLC26A3 | ENSG00000091138 | -4.89597054736947 | -754.177499553705 | 0.330840174402812 | 0.57205899643128 | 24.6878518295235 | 778.865351383229 |
| KLK4 | ENSG00000167749 | -4.78739708526638 | -0.036818346797663 | 0.343745324886879 | 0.724051025161995 | 0.00130138372554361 | 0.0381197305232066 |
| ENSOARG00020005727 | ENSOARG00020005727 | -4.7849627155198 | -0.126265297680224 | 0.883704400412593 | 0.873124015145863 | 0.00447697990330126 | 0.130742277583525 |
| ENSOARG00020002591 | ENSOARG00020002591 | 4.76299285494152 | 0.466048532730646 | 0.678077825251925 | 0.764617399492265 | 0.486036335154241 | 0.0199878024235952 |
| ENSOARG00020014156 | ENSOARG00020014156 | 4.74941696212319 | 4.28441627026882 | 0.716880151074667 | 0.688276805760726 | 4.45875281379534 | 0.174336543526519 |
| S100A8 | ENSG00000143546 | 4.72531298633464 | 1.24619396428815 | 0.763615037471685 | 0.875275348415652 | 1.3678216532036 | 0.12162768891545 |
| KRT36 | ENSG00000126337 | 4.66482018841708 | 0.03647105778499 | 0.773783488764952 | 0.855531537929379 | 0.0376754035582404 | 0.00120434577325037 |

| | | | | | | | |
|---|---|---|---|---|---|---|---|
| GRP | ENSG00000134443 | 4.66367170499564 | 35.9428408560887 | 0.836575551240455 | 0.924344533879189 | 41.3505939564768 | 5.40775310038808 |
| ENSOARG00020021239 | ENSOARG00020021239 | -4.59940498848173 | -2.25380931529806 | 0.931467139560216 | 0.843418264039823 | 0.0950856635040879 | 2.34889497880215 |
| PRSS27 | ENSG00000172382 | 4.54289695514717 | 4.2608055896385 | 0.920812849281677 | 0.717065701003756 | 4.32684874428226 | 0.0660431546437642 |
| ENSOARG00020026387 | ENSOARG00020026387 | -4.48281311685366 | -0.0633855437178787 | 0.607887537161391 | 0.774714330999619 | 0.00279937361951375 | 0.0661849173373925 |
| TMPRSS11D | ENSG00000153802 | 4.47014188018446 | 0.320359227172443 | 0.851097623570003 | 0.759018701265936 | 0.340303618572474 | 0.0199443914000312 |
| ENSOARG00020023153 | ENSOARG00020023153 | -4.46920291737012 | -0.120826576423646 | 0.805968026338844 | 0.655413957691635 | 0.0053391274474405 | 0.126165703871087 |
| RBP2 | ENSG00000114113 | 4.41704619388933 | 1.30153487327226 | 0.558473998831547 | 0.40503014287533 | 1.37119757352733 | 0.0696627002550655 |
| IGFL1 | ENSG00000188293 | 4.35940626188571 | 0.019247284918464 | 0.69833725631857 | 0.816417173255253 | 0.0202812756483838 | 0.00103399072991973 |
| CCK | ENSG00000187094 | 4.31328493529238 | 0.0117975985311752 | 0.518100524620428 | 0.701987852697961 | 0.0124758121098498 | 0.000678213578674543 |
| GYS2 | ENSG00000111713 | 4.3125787380265 | 0.369585906573251 | 0.524253710775918 | 0.685220095218447 | 0.390671053512615 | 0.0210851469393634 |
| S100A12 | ENSG00000163221 | 4.29146525792214 | 2.59106896214359 | 0.760076253812433 | 0.758523901877582 | 2.73791161414256 | 0.146842651998973 |
| ENSOARG00020000383 | ENSOARG00020000383 | 4.26760937076823 | 11.1569115886792 | 0.648259411010195 | 0.600321542206942 | 11.814051924527 | 0.657140335847787 |
| ENSOARG00020011181 | ENSOARG00020011181 | -4.25900027561818 | -0.00225588589752385 | 0.930692106722105 | 0.874560576071345 | 9.1217201147758e-05 | 0.00234710309867161 |
| OXTR | ENSG00000180914 | 4.18480320699374 | 0.547423071976084 | 0.718277754306683 | 0.908907501438242 | 0.897956162912576 | 0.350533090936492 |
| SPINK14 | ENSG00000196800 | -4.1817433953541 | -0.00523798326827404 | 0.987012987012987 | 0.803486387697958 | 5.06417858976757e-05 | 0.00528862505417172 |
| ENSOARG00020023117 | ENSOARG00020023117 | -4.17783768119593 | -0.624688148962628 | 0.698154788809904 | 0.602559400142477 | 0.0342086856612063 | 0.658896834623835 |
| TAC3 | ENSG00000166863 | 4.16692906883266 | 4.6016944536264 | 0.321370647445723 | 0.77216717174452 | 4.89728885568738 | 0.295594402060981 |
| ENSOARG00020025007 | ENSOARG00020025007 | -4.14445638002974 | -8.14973070133269 | 0.572727687120471 | 0.58627115063701 | 0.45598327693389 | 8.60571397826658 |
| LYPD2 | ENSG00000197353 | 4.1303292704533 | 0.302506719475753 | 0.613558304566679 | 0.485074033372085 | 0.322127968800547 | 0.0196212493247939 |
| ENSOARG00020020891 | ENSOARG00020020891 | 4.08819605626998 | 92.8586085438018 | 0.462808121800076 | 0.484643228300231 | 99.1894542548288 | 6.33084571102699 |
| KRT35 | ENSG00000197079 | 4.05551448402891 | 0.758534796303049 | 0.401988086517413 | 0.628226461724914 | 0.809878475410767 | 0.0513436791077181 |
| FSTL5 | ENSG00000168843 | 4.03431199857308 | 0.608568192325194 | 0.503545104226829 | 0.742249302813963 | 0.649788221945426 | 0.0412200296202324 |
| ENSOARG00020010586 | ENSOARG00020010586 | -4.02578428004074 | -0.654413893237148 | 0.879163659198644 | 0.866364404754675 | 0.0415177637863426 | 0.69593165702349 |
| MATN1 | ENSG00000162510 | 4.01864242379543 | 0.0182033427716846 | 0.462937829355593 | 0.616734312399007 | 0.0194741891993515 | 0.00127084642766691 |
| ENSOARG00020005568 | ENSOARG00020005568 | 3.99996950903221 | 8.26166695100927 | 0.556216936306709 | 0.654621653406175 | 8.85229260051368 | 0.590625649504409 |
| ENSOARG00020001244 | ENSOARG00020001244 | -3.9880188404984 | -0.00787557057671479 | 0.795407438558841 | 0.777488005635453 | 0.000490330378307131 | 0.00836590095502192 |
| KRT3 | ENSG00000186442 | 3.95241187819827 | 0.11778235866841 | 0.836574943680045 | 0.712705203734278 | 0.126278816641448 | 0.00849645797303805 |
| MMP12 | ENSG00000262406 | 3.94563871231721 | 2.99494473103739 | 0.529945505351547 | 0.589261116546984 | 3.21901550161837 | 0.224070770580984 |
| ENSOARG00020000427 | ENSOARG00020000427 | 3.93442293948545 | 0.0762251651378047 | 0.768973657959708 | 0.887591724106844 | 0.0821005267021742 | 0.00587536156436946 |
| ATP13A5 | ENSG00000187527 | -3.91530249608752 | -226.767121964689 | 0.610019612038265 | 0.557756088370324 | 15.3212325177328 | 242.088354482421 |
| KRT78 | ENSG00000170423 | 3.84620142858859 | 4.85664628411337 | 0.68743710538597 | 0.510560392168535 | 5.24654141687767 | 0.389895132764298 |
| TMIE | ENSG00000181585 | -3.80542262850698 | -0.501117361694343 | 0.594644543692849 | 0.520508056405506 | 0.0364539900621305 | 0.537571351756473 |
| OLFM4 | ENSG00000102837 | 3.79735004215817 | 4.118938668959 | 0.552708201468705 | 0.653505583247968 | 4.45856250784033 | 0.339623838881322 |
| KRT75 | ENSG00000170454 | 3.78583249517957 | 1.6396310134023 | 0.729285450242233 | 0.7241226686298 | 1.77754891603256 | 0.13791790263026 |
| DKK1 | ENSG00000107984 | -3.78531830978448 | -0.458407213984153 | 0.854845394778049 | 0.52261665445266 | 0.033951079305922 | 0.492358293290075 |
| INHBE | ENSG00000139269 | 3.7554520487291 | 14.0326458083088 | 0.583151242725386 | 0.653289855793245 | 15.2311565181 | 1.19851070979119 |
| IGFBP1 | ENSG00000146678 | 3.75169586700212 | 0.0172064280114894 | 0.654167742730961 | 0.881112440505228 | 0.0183669947090168 | 0.00116056669752738 |
| TTR | ENSG00000118271 | 3.73586741306458 | 0.0328280103927138 | 0.464159129645637 | 0.564666762616246 | 0.0355813833198303 | 0.0027533729271165 |
| ENSOARG00020012854 | ENSOARG00020012854 | 3.73388499719166 | 0.150547505319081 | 0.535012683751303 | 0.45452163732735 | 0.163738578529154 | 0.0131910732100728 |

| | | | | | | | |
|---|---|---|---|---|---|---|---|
| BPIFA3 | ENSG00000131059 | -3.70338876419038 | -0.0665372074337309 | 0.788751878193867 | 0.578375700922563 | 0.00518750493299057 | 0.0717247123667215 |
| ENSOARG00020018060 | ENSOARG00020018060 | -3.61631363751709 | -0.00234932725910245 | 0.96553961435552 | 0.514908196436354 | 0.000163382880576069 | 0.00251271013967852 |
| NMB | ENSG00000197696 | -3.6105884527357 | -0.361948629550567 | 0.421000128689567 | 0.501549653653504 | 0.0302894106364112 | 0.392238040186978 |
| ENSOARG00020005653 | ENSOARG00020005653 | -3.60931247910219 | -0.00842281835341439 | 0.893464683857698 | 0.859496284174818 | 0.000577823043094457 | 0.00900064139650885 |
| ENSOARG00020007834 | ENSOARG00020007834 | 3.60820086544691 | 169.348643046171 | 0.917640829763144 | 0.570628887624536 | 174.100120125419 | 4.75147707924802 |
| APOBEC3Z1 | APOBEC3Z1 | 3.60065605987899 | 0.0818940483603901 | 0.941469453500167 | 0.843196888614979 | 0.0837095119055201 | 0.00181546354513008 |
| AQP9 | ENSG00000103569 | -3.59738285500367 | -86.7124577204827 | 0.696200949984435 | 0.43057366262928 | 7.32754508754471 | 94.0400028080275 |
| NTS | ENSG00000133636 | 3.56316236884929 | 253.658027656422 | 0.50681913545105 | 0.797792179959098 | 278.697144954811 | 25.0391172983889 |
| ENSOARG00020009811 | ENSOARG00020009811 | 3.56249925596615 | 0.558084808281715 | 0.831713052585341 | 0.85159934279402 | 0.612167529601945 | 0.0540827213202299 |
| GREM1 | ENSG00000282046 | 3.55628746191299 | 3.35950016879503 | 0.647308286433341 | 0.525331956299854 | 3.70107558490097 | 0.341575416105935 |
| ENSOARG00020012216 | ENSOARG00020012216 | -3.49032633160166 | -18.1010250730951 | 0.601089691235662 | 0.554103952875788 | 1.68238105983719 | 19.7834061329323 |
| SERPINB12 | ENSG00000166634 | 3.47420778507579 | 0.192089898033723 | 0.556225483769819 | 0.56136095265624 | 0.212634844490066 | 0.0205449464563438 |
| ENSOARG00020001753 | ENSOARG00020001753 | -3.43537988949491 | -0.221251173376968 | 0.698080404451732 | 0.608363666918682 | 0.0210162606866806 | 0.242267434063648 |
| ENSOARG00020018226 | ENSOARG00020018226 | -3.40717661733745 | -0.0532386590988594 | 0.749107609830539 | 0.541064883342557 | 0.00522088185396742 | 0.0584595409528269 |
| ENSOARG00020001098 | ENSOARG00020001098 | -3.40344001263939 | -0.0342357893661368 | 0.830611709397833 | 0.730729588621107 | 0.00328916821087223 | 0.037524957577009 |
| PHLDA2 | ENSG00000274538 | 3.39106716508031 | 0.325306474319888 | 0.557379592939645 | 0.477502454620655 | 0.362979900512606 | 0.0376734261927181 |
| ENSOARG00020007580 | ENSOARG00020007580 | 3.38023399394846 | 14.6884611380242 | 0.916202274448986 | 0.547370017050209 | 15.1751004962149 | 0.486639358190693 |
| ENSOARG00020008854 | ENSOARG00020008854 | -3.32043361197932 | -14.3420626855636 | 0.613757392231996 | 0.504421717314006 | 1.51535317788897 | 15.8574158634526 |
| UPK2 | ENSG00000110375 | 3.31434756135508 | 0.00313727307173313 | 0.527111993891946 | 0.794120860470912 | 0.00352460930929406 | 0.000387336237560927 |
| MMP7 | ENSG00000137673 | 3.29494686122149 | 3.01760775813671 | 0.743589577048928 | 0.769719728775387 | 3.92029956663954 | 0.902691808502828 |
| GRXCR1 | ENSG00000215203 | 3.26416933728337 | 0.79133409904993 | 0.48311096262132 | 0.827128452731347 | 0.874049535012958 | 0.0827154359630285 |
| GREB1 | ENSG00000196208 | 3.26232532203146 | 976.398750184738 | 0.309089605914334 | 0.440305795982338 | 1099.41139428492 | 123.012644100181 |
| ENSOARG00020008814 | ENSOARG00020008814 | -3.26060519749211 | -0.244046050661095 | 0.755270394178281 | 0.548891648029976 | 0.0269150977481888 | 0.270961148409283 |
| GJB6 | ENSG00000121742 | 3.25789093422036 | 0.00483369541035877 | 0.733228680166457 | 0.827009795359644 | 0.00533650100566402 | 0.000502805595305254 |
| ENSOARG00020000477 | ENSOARG00020000477 | 3.24317205802819 | 0.0354886233817352 | 0.824143541036406 | 0.670801312676388 | 0.038286804915271 | 0.00279818153353582 |

**Table S5:** Genes exclusive to bovine cervix in the follicular and luteal phase

| Gene | ID | **Log Fold Change** | **Dif Means** | gini_1 | gini_2 | Mean1 | Mean2 | estrus phase |
|---|---|---|---|---|---|---|---|---|
| BPIFA2B | ENSBTAG00000019752 | 8.01169488659356 | 0.627058853372568 | 0.495585367076419 | NA | 0.627058853372568 | 0 | Follicular |
| LOC112446672 | ENSBTAG00000052798 | 5.97028153990243 | 0.422175917456848 | 0.568501932138137 | NA | 0.422175917456848 | 0 | Follicular |
| RDH16 | ENSBTAG00000001392 | -5.30278893747946 | -0.211260754246596 | NA | 0.22153142686123 | 0 | 0.211260754246596 | Luteal |
| C29H11orf86 | ENSBTAG00000022246 | 5.11490131443356 | 0.901634401094783 | 0.466235913936583 | NA | 0.901634401094783 | 0 | Follicular |
| ENSBTAG00000048830 | ENSBTAG00000048830 | 4.80228490133332 | | 0.423799663506785 | NA | 0.0270053431419432 | 0 | Follicular |
| TMEM229A | ENSBTAG00000049382 | 4.66450793972288 | 0.192363626559942 | 0.441941247442293 | NA | 0.192363626559942 | 0 | Follicular |
| ENSBTAG00000005324 | ENSBTAG00000005324 | 4.58816104654107 | 2.82899586567018 | 0.572272110442103 | NA | 2.82899586567018 | 0 | Follicular |
| BSP3 | ENSBTAG00000003886 | -4.55507383233905 | -0.640788200719888 | NA | 0.358662173213077 | 0 | 0.640788200719888 | Luteal |
| LOC507527 | ENSBTAG00000013507 | 4.33049803539077 | 0.00796028178064769 | 0.562107108912761 | NA | 0.00796028178064769 | 0 | Follicular |
| NPVF | ENSBTAG00000019447 | 4.19515510583535 | 0.0412841844376611 | 0.459296453953889 | NA | 0.0412841844376611 | 0 | Follicular |
| ENSBTAG00000052976 | ENSBTAG00000052976 | -4.14806392999975 | -0.135683077496103 | NA | 0.268062655301592 | 0 | 0.135683077496103 | Luteal |
| ENSBTAG00000012533 | ENSBTAG00000012533 | 4.12932092887349 | 0.00941592543141604 | 0.62449535675828 | NA | 0.00941592543141604 | 0 | Follicular |
| ENSBTAG00000049959 | ENSBTAG00000049959 | 3.72021199993998 | 0.00705352483514716 | 0.519300181975886 | NA | 0.00705352483514716 | 0 | Follicular |
| LOC520402 | ENSBTAG00000040320 | -3.39489882153536 | -0.0364767605915812 | NA | 0.393006256335909 | 0 | 0.0364767605915812 | Luteal |
| KCNC2 | ENSBTAG00000054123 | 3.36487699970871 | 0.0813137372751472 | 0.615161872409253 | NA | 0.0813137372751472 | 0 | Follicular |
| MYL10 | ENSBTAG00000026273 | 3.34370470861356 | 0.121832086532478 | 0.620855813182252 | NA | 0.121832086532478 | 0 | Follicular |
| R3HDML | ENSBTAG00000013220 | 3.1956982497018 | 0.0211369823488593 | 0.614225023129965 | NA | 0.0211369823488593 | 0 | Follicular |
| LOC523389 | ENSBTAG00000053227 | 3.1156670054286 | 0.0553082185424062 | 0.440599361865036 | NA | 0.0553082185424062 | 0 | Follicular |
| ENSBTAG00000040248 | ENSBTAG00000040248 | 2.98069741015961 | 0.0126126783125132 | 0.422518946319597 | NA | 0.0126126783125132 | 0 | Follicular |
| MC4R | ENSBTAG00000019676 | 2.77976167767441 | 0.181892877426001 | 0.647587296803692 | NA | 0.181892877426001 | 0 | Follicular |
| ENSBTAG00000054106 | ENSBTAG00000054106 | -2.77866487828753 | -0.032596894581462 | NA | 141339065763543 | 0 | 0.032596894581462 | Luteal |
| ENSBTAG00000053691 | ENSBTAG00000053691 | 2.67531653468796 | 0.00638802558021577 | 0.329641811491069 | NA | 0.00638802558021577 | 0 | Follicular |
| ENSBTAG00000054917 | ENSBTAG00000054917 | -2.54230941311803 | -0.0169661005586573 | NA | 0.23921970533918 | 0 | 0.0169661005586573 | Luteal |
